# Neurotranscriptomic profiling of DWV-infected honey bee foragers with different cognitive abilities

**DOI:** 10.1101/2025.05.23.655587

**Authors:** Simon Loughran, Lauren Dingle, Alan S. Bowman, Fabio Manfredini

**Affiliations:** School of Biological Sciences, University of Aberdeen, AB24 3UL Aberdeen, UK

**Author notes:** Correspondence (SL); (FM). (S.L.); (L.D.); (A.S.B); (F.M.).

**Keywords:** deformed wing virus, *Apis mellifera*, RNA-seq, transcriptome, Gene Regulatory Networks, mushroom bodies, brain, behaviour, Proboscis Extension Reflex, Associative Learning

## Abstract

Honey bees (*Apis mellifera*) provide important ecosystem services to both natural and human-managed environments, but are increasingly threatened by a variety of pathogens, the most common of which is deformed wing virus (DWV). DWV is known to replicate in the honey bee brain and has been documented as both improving and impairing olfactory learning and memory. We examined the transcriptomic response of the honey bee mushroom bodies—an area of the insect brain associated with higher cognitive functions—in bees with naturally occurring DWV infections who varied in their ability to perform an associative learning task. RNA-seq analysis detected increased expression of genes involved in the immune response, including important antimicrobial peptides (AMPs) such as *hymenoptaecin, apidaecin*, and *abaecin*, and the downreguation of *lysozyme*, *PPO*, and other genes associated with responses to a range of stressors. Additionally, gene ontology (GO) enrichment analysis revealed overrepresentation of key biological processes which form part of the immune response. We also noted significant differential expression of long non-coding RNAs (lncRNAs) presumed to be acting in a regulatory manner, and used these lncRNAs to construct gene regulatory networks (GRNs). Strikingly, in contrast to previous studies on bees with artificially-induced infections that have examined viral loads in the abdomen and non-specific areas of the brain, no correlation between DWV load in the mushroom bodies and cognitive function was noted. This highlights the complexity of host-pathogen interactions in honey bee neural tissues and the benefits of a spatially-refined approach to brain transcriptomics in naturally-occurring infections.

## 1. Introduction

Insect pollinators, like many bee species, provide important ecosystem services to both natural and human-managed environments, favouring the transfer of pollen from male to female flower parts with their incessant foraging activity. Foraging requires not only the high energy expenditure associated with flight—a metabolically intensive behaviour [1,2]—but also a sophisticated suite of cognitive skills, including the ability to orient, navigate, memorise and learn [3,4]. It is well established that a range of stressors can take a significant toll on the foraging behaviour of many bee species [5], either by affecting their energy levels and metabolism [6], by altering their floral preferences [7], or by directly compromising their cognitive abilities [5,8]. For example, land-use and climate change, and resultant habitat loss and fragmentation, are associated with a reduction in the richness, diversity, and abundance of wild bees, smaller body sizes, and a decline in pollination services [9,10]. In honey bees, exposure to sublethal doses of many insecticides can affect learning and memory [11,12], including specific effects on the mushroom bodies [13,14], while parasites and pathogens can impact multiple traits, including foraging activity and age of onset of foraging, social interactions, learning and memory, and motor behaviour [8]. Honey bee colonies are typically simultaneously exposed to multiple stressors resulting in complex interactions and antagonistic and synergistic effects [15,16], and a compromised immune system, which can exacerbate the impact of each stressor [17]. It is noteworthy that many of the observed effects are similar for different stressors, suggesting similar mechanisms of actions within target tissues such as the brain. An integrated honey bee stress pathway has been hypothesised [18]; however, this model focuses on a general stress response to acute and short-term stressors, and the mechanisms by which key stressors alter brain functions over a longer period of time to produce the observed behavioural changes remain largely uncharacterised.

The majority of viruses infecting honey bees are single-stranded RNA viruses, icosahedral in shape, 30 nm in size, and do not cause symptoms in individuals that are infected [19–21]. Both horizontal and vertical transmission have been observed, and infection and presence of disease can vary according to life stage and season [19,20,22]. Of these, deformed wing virus (DWV) is the most widespread and is recognised as a major honey bee stressor and contributor to colony demise [23,24]. When transmitted by the ectoparasite *Varroa destructor*, which feeds on the haemolymph of honey bee pre-imaginal stages [25–27], DWV reaches extremely high levels, capable of producing abnormalities at the morphological and physiological levels such as deformed wings and bloated abdomens, and can also lead to early mortality [28]. However, DWV can also be transmitted orally *via* trophallaxis or shared resources [29], and venerally and vertically [30]. Oral transmission typically results in asymptomatic infections or milder symptoms, such as an early transition from nursing to foraging and impaired learning [31]. Disruption of foraging behaviour itself, such as a reduction in flight distance and duration, can also occur [32]. It is generally assumed that asymptomatic infections are better tolerated by honey bee colonies, but preliminary evidence has shown that even asymptomatic infections could have a significant toll on colony fitness and survival [33,34], warranting further investigation. Furthermore, the mechanisms by which DWV produces the reported behavioural changes are largely unknown and their study could contribute to the broader field of the neurobiology of insects.

DWV has been detected in several regions of the honey bee brain (or neuropils) such as the optic lobes, antennal lobes, and mushroom bodies [35]. The presence of the virus in these regions strongly supports the hypothesis that infection may impact various sensory activities essential for a bee forager to perform effectively, such as visually identifying rewarding flowers and detecting the scent or sugar concentration of nectar. DWV localisation in mushroom bodies is particularly noteworthy since this region of the insect brain is home to important cognitive functions such as learning and memory, which are fundamental for foragers to recall which flowers they have visited, and which are most rewarding. Intriguingly, previous research has shown that not only the genomic strand but also the replicating strand of DWV can be detected in the mushroom bodies of infected bees [35], supporting the hypothesis that this virus might interfere directly with neural functions associated with foraging behaviour.

Many studies have examined the impact of viral infections on abdomen or whole body transcriptomes of honey bees [36], and analysis of brain transcriptomes has become more common in recent years [37]. Pizzorno et al. [38] have recently specifically explored differentially expressed genes in isolated brain tissue from bees exposed to abdominal injections of DWV inoculum. The study reported the overexpression of genes associated with immune response, and given the documented presence of DWV in the mushroom bodies, we hypothesise that immune-related genes should also be present more specifically in this area.

The link between DWV and honey bee behaviour has been explored only partially. Previous research indicates that DWV-infected honey bees perform poorly in classic learning and memory tasks, as evaluated with the Proboscis Extension Reflex (PER) [39,40], a typical association paradigm that is used to assess cognitive abilities in a fully controlled experimental setup. However, these studies typically introduce the virus into adult worker bees using artificial methods—either by injection with a syringe to mimic natural *Varroa*-mediated transmission, or by feeding to replicate natural oral routes of infection. As such, some important steps associated with the pathogenicity and tissue tropism of DWV in the pre-imaginal stages of the host are bypassed with these approaches. This is a key moment in the interaction between host and pathogen, since the majority of natural infections begin precisely at the pre-imaginal stage and could have significant consequences for later developmental stages. Most importantly, the honey bee neural system is still not fully formed at the pre-imaginal stages and only reaches full maturity in adult bees [41]. Intriguingly, a recent study by Szymański et al. [42] investigated the effect on cognitive abilities of natural DWV infections in the mushroom bodies of honey bee foragers and, in contrast to what has been reported in previous research, detected a subtle positive effect of the virus on reversal learning—but no effect in a simple associative learning task. This demonstrates a need for a focus on natural DWV infections, in order to provide a more comprehensive account of the interplay between this virus—in particular when detectable in the brain—and honey bee cognitive abilities, highlighting potential trade-offs between brain functions and immune responses that are triggered by viral infections [38,43].

Here we focused on the analysis of honey bee foragers displaying natural infections with DWV loads ranging from low to high (but still lacking the deformities that would render a worker bee unviable as a forager). We assessed the cognitive abilities of these bees using a simple associative PER assessment combined with intermediate-memory retention, and selected bees that performed very well in the task (Good Learners) and bees that failed completely (Poor Learners). We then quantified with RT-qPCR the exact number of DWV copies (or genome equivalent) in the mushroom bodies of bees belonging to the two groups and performed a transcriptomic profiling of the mushroom bodies of these bees, with the aim of detecting the most-significant gene expression profiles associated with cognitive performance and DWV infection.

## 1. Materials and Methods

### 2.1. Honey Bee Samples

The honey bee samples used in this study were obtained from two colonies located in two apiaries near Aberdeen, Scotland: Cruickshank Botanical Garden, on the University of Aberdeen Kings College campus (Grid Reference NJ936085), and in Newburgh (NJ998260). Colonies in Newburgh (Colony A) receive minimal *Varroa* treatments and therefore suffer from relatively high *Varroa* and DWV levels [44]. In contrast, colonies in Cruickshank Botanical Garden (Colony B) receive standard treatments against the parasitic mite *Varroa destructor*, and consequently have low Varroa infection rates and low DWV levels. The two apiaries were specifically targeted to obtain honey bee samples with expected significant variation in viral loads. Honey bee foragers were collected in the summer of 2020 between 9.00 and 10.00 at the colony entrance when returning from a foraging trip. The bees were transported individually in petri dishes, and once in the lab were immobilised on ice and harnessed for an absolute conditioning assay *via* PER (Proboscis Extension Reflex).

### 2.2. PER Conditioning Assays

This set of experiments was performed to quantify the learning performance of individual bees and thereby allocate each to one of two distinct behavioural groups: Good Learners and Poor Learners. Harnessed bees were initially fed with 3–5 μL of 30% (w/w) sucrose solution and desensitised to water by touching the antennae with a soaked toothpick. They were then held in the harnesses for one hour before the start of the assays. The general PER conditioning protocol followed the approach of [45], but with only one scent being used and no reversal performed. The bees were conditioned to respond to citral (Sigma-Aldrich, Saint Louis, Missouri, USA) through a 5-time exposure to the odour with reinforcement by touching the antennae with a 30% (w/w) sucrose solution. 5 μL citral was pipetted onto a piece of filter paper, which was then inserted into a 20 mL syringe. The bees were exposed to citral by expressing the full volume of air from the syringe around 0.5 cm away from the antennae. Foragers that exhibited PER to the first citral exposure were discarded. If no PER was recorded for the odour itself, the stimulus was reinforced with a 30% (w/w) sucrose solution. Individuals not responding to sucrose were discarded before further testing.

After pre-conditioning, but on the same day as capture, bees were re-fed and exposed to the same odour three times without reinforcement—this was repeated the following day for a total of six tests. Samples were then allocated to the two behavioural groups according to PER response in the non-reinforced trials as follows: Poor Learners = no response in any of the trials; Good Learners = PER response in all 6 trials. Thereafter, bees were frozen in a -80°C freezer and stored there until later processing for molecular work.

### 2.3. Processing of Samples for Molecular Work

Total RNA was isolated from the mushroom bodies of individual bees for two purposes: 1) to quantify DWV loads in this tissue and 2) to perform transcriptomic profiling with RNA-seq. Mushroom bodies were dissected on dry ice under a stereomicroscope following [42], and homogenised in a Tissue Lyser II device (Qiagen, Hilden, Germany) using 1 ml TRIzol and 2.3 mm zirconia beads (Thistle Scientific, Glasgow, UK). We then used a standard TRIzol protocol to obtain total RNA that was eluted in 20 μl of nuclease-free water. These RNA samples were used for the quantification of DWV loads with an RT-qPCR assay: 300 ng of RNA was converted into cDNA using the iScriptTM cDNA Synthesis Kit (BioRad Laboratories Inc., Hercules, California, USA) and following the manufacturer’s protocol. cDNA samples (300 ng) were used as templates to run RT-qPCR reactions with primers capable of detecting the two most common strains of DWV [46]. DWV loads were quantified as genome equivalents (GE) using calibration curves obtained from known quantities of DWV GE inserted into plasmids.

Bees were allocated to two groups as follows: high DWV—when mushroom bodies harboured more than log_10_ 5 DWV GE overall, and low DWV—when viral loads were lower than or equal to log_10_ 4 DWV GE overall. By combining the behavioural grouping with the grouping resulting from DWV analyses, we were able to obtain a total of 33 bee mushroom body samples (Table 1) that underwent RNA-seq profiling. These samples were processed via a clean-up and concentration step that also included the removal of residual genomic DNA (Zymo Research Corp., Irvine, California, USA) and thereafter they were sent to the Centre for Genome Enabled Biology and Medicine at the University of Aberdeen for library preparation and sequencing.

**Table 1.** Summary of the 33 mushroom body samples used in this study.

| Sample ID | Viral Load (log<sub>10</sub>) | Viral Load <sup>1</sup> | Performance <sup>2</sup> | Colony <sup>3</sup> |
| --- | --- | --- | --- | --- |
| 1 | 2.95 | low | poor | B |
| 2 | 2.51 | low | poor | B |
| 3 | 2.59 | low | poor | A |
| 4 | 2.62 | low | poor | A |
| 5 | 3.96 | low | good | A |
| 6 | 2.49 | low | good | A |
| 7 | 4.01 | low | good | B |
| 9 | 3.56 | low | poor | B |
| 10 | 5.18 | high | poor | B |
| 11 | 2.68 | low | poor | A |
| 12 | 10.35 | high | poor | A |
| 13 | 4.05 | low | good | B |
| 14 | 9.83 | high | good | B |
| 15 | 2.75 | low | good | A |
| 16 | 9.25 | high | good | A |
| 17 | 2.49 | low | poor | B |
| 18 | 2.81 | low | poor | A |
| 19 | 3.02 | low | good | A |
| 20 | 2.38 | low | good | B |
| 21 | 9.36 | high | poor | A |
| 22 | 6.71 | high | poor | B |
| 23 | 6.47 | high | good | A |
| 24 | 9.02 | high | good | B |
| 25 | 2.52 | low | poor | B |
| 26 | 2.88 | low | poor | A |
| 27 | 2.98 | low | good | A |
| 28 | 2.44 | low | good | B |
| 29 | 10.35 | high | poor | A |
| 30 | 5.14 | high | poor | B |
| 31 | 5.28 | high | good | A |
| 32 | 6.38 | high | good | B |
| 33 | 5.26 | high | good | B |
| 34 | 5.35 | high | good | B |

<sup>1</sup> high = DWV GE $\geq \log_{10} 5$ ; low = DWV GE $\leq \log_{10} 4$ (total quantity of virus in the mushroom bodies). <sup>2</sup> poor = Poor Learners who failed all trials; good = Good Learners who passed all trials. <sup>3</sup> Colony A —bees collected from Newburgh; Colony B—bees collected from Cruickshank.

Library prep was performed using a TruSeq Stranded mRNA-seq Kit: the 33 samples were split into two batches (balanced by treatments and other major variables) and indexed for Illumina sequencing. Sequencing was carried out on a NextSeq500 v2.5 platform using Illumina High Output v2.5 75 cycle flowcell and reagents. The output produced a total of 874.2M single-end 75 bp reads—on average 23.2M per sample.

### 2.4. Processing of RNA-seq data

Raw reads were trimmed using Trim Galore! v0.6.4 [47]. Illumina adapter sequences were removed using a stringency setting of 3; ends including bases with a phred score less than 30 were removed; and reads shorter than 30 base pairs were discarded. Read quality was then assessed with FastQC v0.11.8 [48]. There was no evidence of adaptor contamination, but six samples—five of which carried high viral loads—included overrepresented sequences, lower-than-average alignments, and slightly atypical per-sequence GC content plots, suggesting contamination. We extracted these overrepresented sequences and used BLAST (Altschul et al. 1990) to find matches in the NCBI standard nucleotide sequence databases, which revealed a close match to DWV genomes. The DWV reads were identified using HISAT2 v2.1.0 [49]: reads from each sample were aligned to DWV reference genomes—a recombinant strain (NCBI accession: MN538210) [50] and a DWV-B strain (NCBI accession: MN565038) [51]. Options were set so that separate files were generated for aligned reads and unaligned reads. Table 2 summarises the effects of removing the DWV alignments. The unaligned reads for these six samples then served as “decontaminated” reads to replace the originals. These and the remaining 27 sample reads were aligned to the *A. mellifera* genome assembly (Amel_HAv3.1; NCBI Assembly: GCF_003254395.2). FeatureCounts (from Subread v1.6.2) [52] was used to obtain raw read counts used for analysis.

**Table 2.** The six mushroom body samples whose reads included overrepresented sequences. Reads aligning to the DWV genome were removed and the percentage of overrepresented sequence before and after removal is shown. The expected source of any remaining overrepresented sequences (according to a BLAST search) is also noted.

| Sample ID | Viral Load <sup>1</sup> | Before (%) | After (%) | Source |
| --- | --- | --- | --- | --- |
| 12 | High | 9.1 | 0 |  |
| 14 | High | 11.1 | 0 |  |
| 16 | High | 1.0 | 0 |  |
| 20 | Low | 0.2 | 0 |  |
| 21 | High | 0.6 | 0.6 | Ribosomal RNA |
| 29 | High | 2.2 | 0 |  |

<sup>1</sup> high = DWV GE $\geq \log_{10} 5$ ; low = DWV GE $\leq \log_{10} 4$ .

### 2.5. Exploration and analyses

We performed exploratory analyses on the RNA-Seq dataset with principal component analyses (PCA) to investigate variations in patterns of gene expression due to colony location, learning performance and viral load. In order to reduce noise, enhance biological signal, and to improve visualisation and clustering [53–55], analyses were conducted on the normalised gene expression counts for the 500 most-variable of the 12,332 genes from the 33 mushroom body samples. Raw counts were normalised using the median of ratios method and applying a variance stabilising transformation [56], and a Pearson correlation coefficient matrix was created for the pairwise comparison of samples. Noting greater clustering amongst Colony A samples, we quantified clustering strength by measuring silhouette width [57] and the within-colony Euclidean distances in PCA space.

We carried out differential gene expression analysis using DESeq2 v1.40.2 [55]. To ensure only genes with sufficiently large read counts across samples were used in the analysis we filtered the list of genes according to expression level by using the filterByExpr function from edgeR v 3.42.4 [58], and then used DESeq2 to reveal a list of differentially expressed genes with a Benjamini-Hochberg-adjusted p-value < 0.05 and an absolute log_2_ fold change in expression of one or more (|log₂FC| > 1). For each differentially expressed gene we also examined the Spearman’s correlation coefficient (SCC) between absolute viral load (see Table 1) and the normalised gene expression level. We also used Bioconductor’s EnrichmentBrowser v 2.30.2 [59] to assemble gene sets from the expression data using over-representation analysis (ORA) [60] and tested for enrichment of functional gene sets as defined in gene ontology (GO) [61] and Kyoto Encyclopedia of Genes and Genomes (KEGG) [62] pathway annotations. Finally, we used BioNero’s [45] exp2grn function to explore gene regulatory networks (GRNs), using differentially expressed lncRNAs as potential regulators of all genes.

## 3. Results

### 3.1. Overall patterns of gene expression in honeybee mushroom bodies

Principal Component Analyses showed that samples clearly separated on both PC1 and PC2 according to colony of origin (Figure 1). There was little clustering due to behavioural test score, suggesting no clear effect of learning performance on global gene expression. A Scree plot is included in Figure S1.

**Figure 1.**
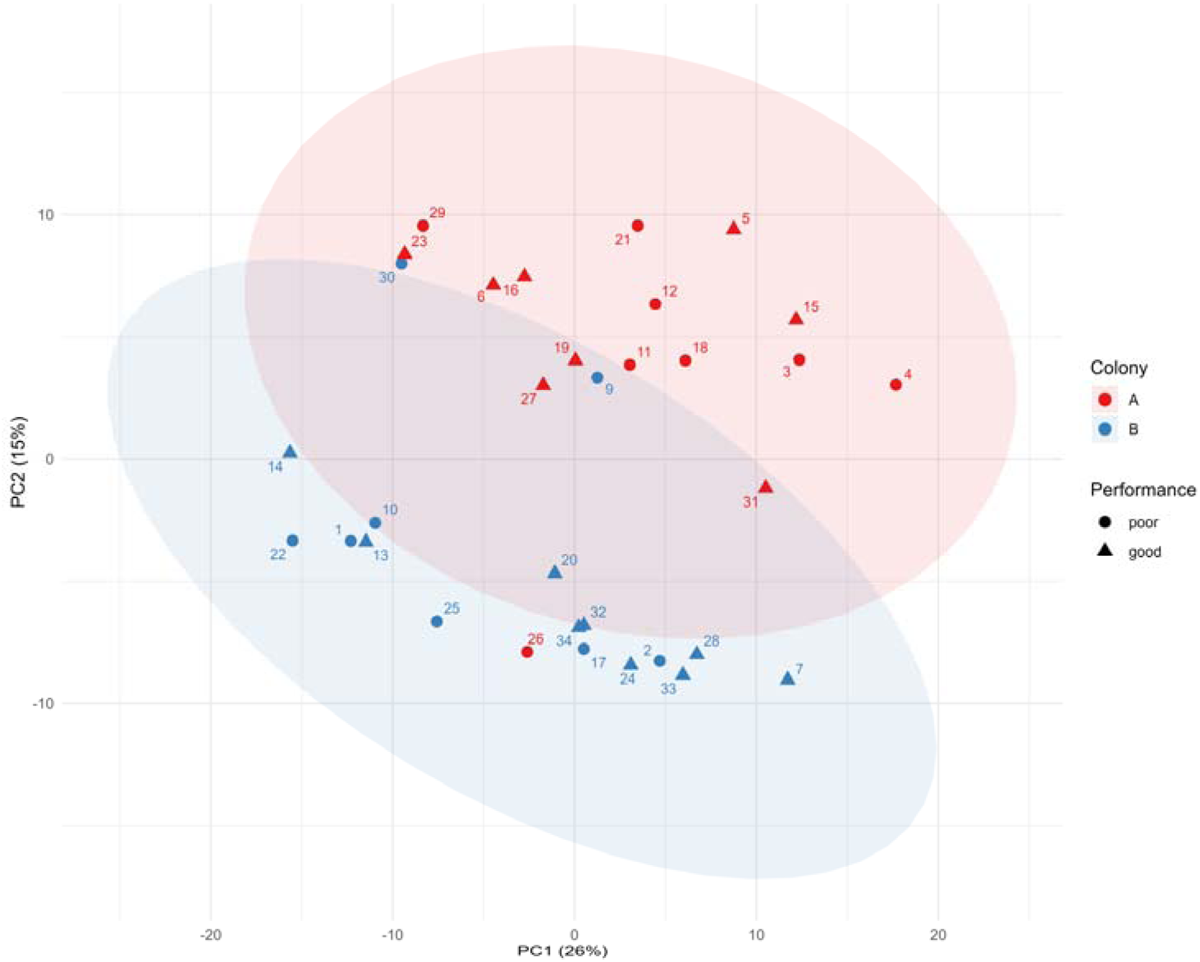
Principal component analysis showing clustering of the 33 mushroom body samples according to colony of origin, and absence of clustering according to performance in behavioural tests. PCA was performed using the normalised gene expression counts for the 500 most-variable genes expressed in the samples. Each point represents one of the 33 samples and is colour-coded to indicate colony of origin, and shaped to indicate performance. 95% confidence ellipses are superimposed for colony of origin. PCs 1 and 2 together account for 41% of variation.

### 3.2. Transcriptomic profiling of different groups of learners

Given that the main goal of this study was to assess the impact of mushroom body viral infection on learning performance—and to explore whether this is reflected at the transcriptomic level—we created subsets of good and poor learners, and within each subset compared gene expression levels of individuals with high and low viral loads. These more-nuanced analyses gave us greater power to identify the subtle effects of viral infection on learning performance. The analysis on Poor Learners revealed a set of 58 genes that were significantly differentially expressed (Benjamini-Hochberg-adjusted *p* < 0.05). Of these, 15 genes were expressed with an absolute log_2_ fold change of one or more (|log₂FC| > 1) (Figure 2; Table 3). 80% of these genes were expressed at lower levels in Poor Learners when viral infection was high.

GO analysis revealed enrichment (Bonferroni-adjusted *p* < 0.01) of key groups: GO:0007601/visual perception (LOC726228, *Lop2, Lop1*), GO:0045087/innate immune response, and GO:0002376/immune system process (*Apid1*, LOC406142, LOC406144); there was also enrichment of GO:0004930 and GO:0007186, G protein-coupled receptor signaling pathways. KEGG pathway analysis revealed pathways associated with phototransduction and, neuroactive ligand-receptor interaction, but neither of these was significant following the Bonferroni correction (see Table S1). When comparing bees with high vs. low viral load within the subset of Good Learners no genes were significantly expressed.

**Figure 2.**
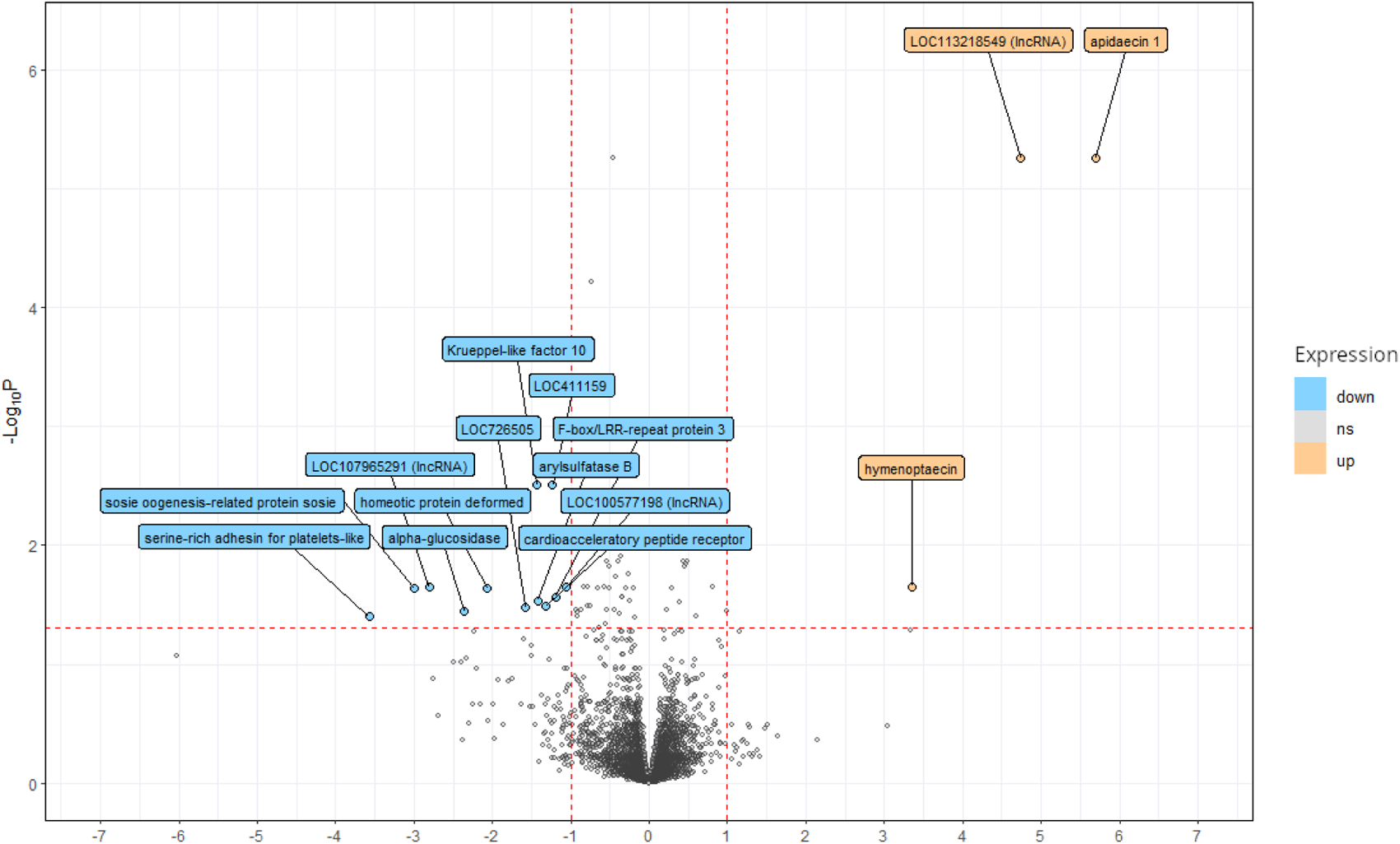
Volcano plot showing significantly expressed genes in Poor Learner bee mushroom bodies (Benjamini-Hochberg-adjusted *p* < 0.05, horizontal red dotted line) with an absolute log_2_ fold change of 1 or more (|log₂FC| > 1) (vertical red dotted lines). Genes in orange are more highly expressed in Poor Learners with high viral loads (≥ log_10_ 5 DWV GE), while genes in blue are expressed at lower level in Poor Learners with high viral loads. Both measurements reported with reference to Poor Learners with low viral loads (≤ log_10_ 4 DWV GE).

**Table 3.** List of genes differentially expressed in mushroom bodies of Poor Learners when comparing individuals with high and low viral loads (DWV ≥ log_10_ 5 and ≤ log_10_ 4, repectively).

| Regulation <sup>1</sup> | OGSv3.2 ID <sup>2</sup> | OGSv1.x ID <sup>3</sup> | Gene ID <sup>4</sup> | Gene Description | Log<sub>2</sub> FC <sup>5</sup> | *p* <sup>6</sup> | Rho <sup>7</sup> |
| --- | --- | --- | --- | --- | --- | --- | --- |
| Up | GB46236, GB47546, GB51306 | GB13473, GB17354, GB17782 | 406140, *Apid1* | apidaecin 1 | 5.69 | $5.63 \times 10^{-6}$ | 0.58 |
| | | | LOC113218549 | non-coding RNA | 4.74 | $5.63 \times 10^{-6}$ | 0.34 |
| | GB1223 | GB17538 | LOC406142 | hymenoptaecin | 3.35 | $2.22 \times 10^{-2}$ | 0.27 |
| Down | GB48020 | GB13606 | LOC102655185 | serine-rich adhesin for platelets-like | 3.57 | $3.93 \times 10^{-2}$ | 0.65 |
| | GB42311 | GB14425 | LOC102654257 | sosie oogenesis-related protein sosie | 2.99 | $2.31 \times 10^{-2}$ | 0.65 |
| | | | LOC107965291 | non-coding RNA | 2.81 | $2.22 \times 10^{-2}$ | 0.75 |
| | GB43247 | GB19017 | LOC406131, *Hbg3* | alpha-glucosidase | 2.36 | $3.58 \times 10^{-2}$ | 0.61 |
| | GB51299 | GB13409 | LOC724252, *Dfd* | homeotic protein deformed | 2.06 | $2.32 \times 10^{-2}$ | 0.69 |
| | GB41033, GB41034 | GB11097, GB14306 | LOC726505 | uncharacterized protein coding | 1.58 | $3.34 \times 10^{-2}$ | 0.77 |
| | GB45040 | GB10114 | LOC410326 | Krueppel-like factor 10 | 1.43 | $3.11 \times 10^{-3}$ | 0.61 |
| | GB40771 | GB13235 | LOC412829 | arylsulfatase B | 1.42 | $2.92 \times 10^{-2}$ | 0.55 |
| | GB54316 | GB19289, GB30535 | LOC726935 | Crustacean cardioactive peptide receptor | 1.32 | $3.27 \times 10^{-2}$ | 0.55 |
| | GB52955 | GB11664 | LOC411159 | uncharacterized protein coding | 1.23 | $3.11 \times 10^{-3}$ | 0.76 |
| | GB49376 | GB19733 | LOC724642 | F-box/LRR-repeat protein 3 | 1.20 | $2.71 \times 10^{-2}$ | 0.79 |
| | GB55973 | | LOC100577198 | uncharacterized protein coding | 1.05 | $2.22 \times 10^{-2}$ | 0.39 |

<sup>1</sup> Up = more highly expressed in Poor Learners with high viral loads; Down = lower expression in Poor Learners with high viral loads. <sup>2</sup> Gene IDs as specified in Official Gene Set (OGS) version 3.2 [63]. <sup>3</sup> Gene IDs as specified in Official Gene Set (OGS) version 1.x [64]. <sup>4</sup> NCBI Gene ID (https://www.ncbi.nlm.nih.gov/gene/). <sup>5</sup> Log<sub>2</sub> FC = absolute difference in gene expression expressed as fold changes on a log<sub>2</sub> scale ( $|\log_2 FC|$ ). <sup>6</sup> Bonferroni-adjusted p-value. <sup>7</sup> Rho = correlation coefficient between normalised read counts and DWV loads.

### 3.3. Transcriptomic profiling in response to viral infection

PCA revealed that colony is an important factor driving global patterns of gene expression in this study (Figure 1). It is of note that Colony A included four samples with very high viral loads (> 9 log_10_), compared to only two such samples from Colony B, reflecting the typically higher prevalence of DWV in the apiary incorporating Colony A [46]. In addition, a heatmap analysis suggested greater clustering for Colony A compared to Colony B (Figure 3) and this pattern was confirmed with cluster analysis: Colony A samples exhibited significantly tighter transcriptomic clustering relative to Colony B, as indicated by a higher average silhouette width (0.146 vs 0.010, *p* = 1.57 × 10⁻⁶, Wilcoxon test) and a lower mean within-group distance in PCA space (15.2 vs 21.7, *p* = 0.00084, Wilcoxon test), calculated using the first three principal components, which account for 49.7% of total variance. These results confirm a greater degree of transcriptomic similarity among Colony A bees.

**Figure 3.**
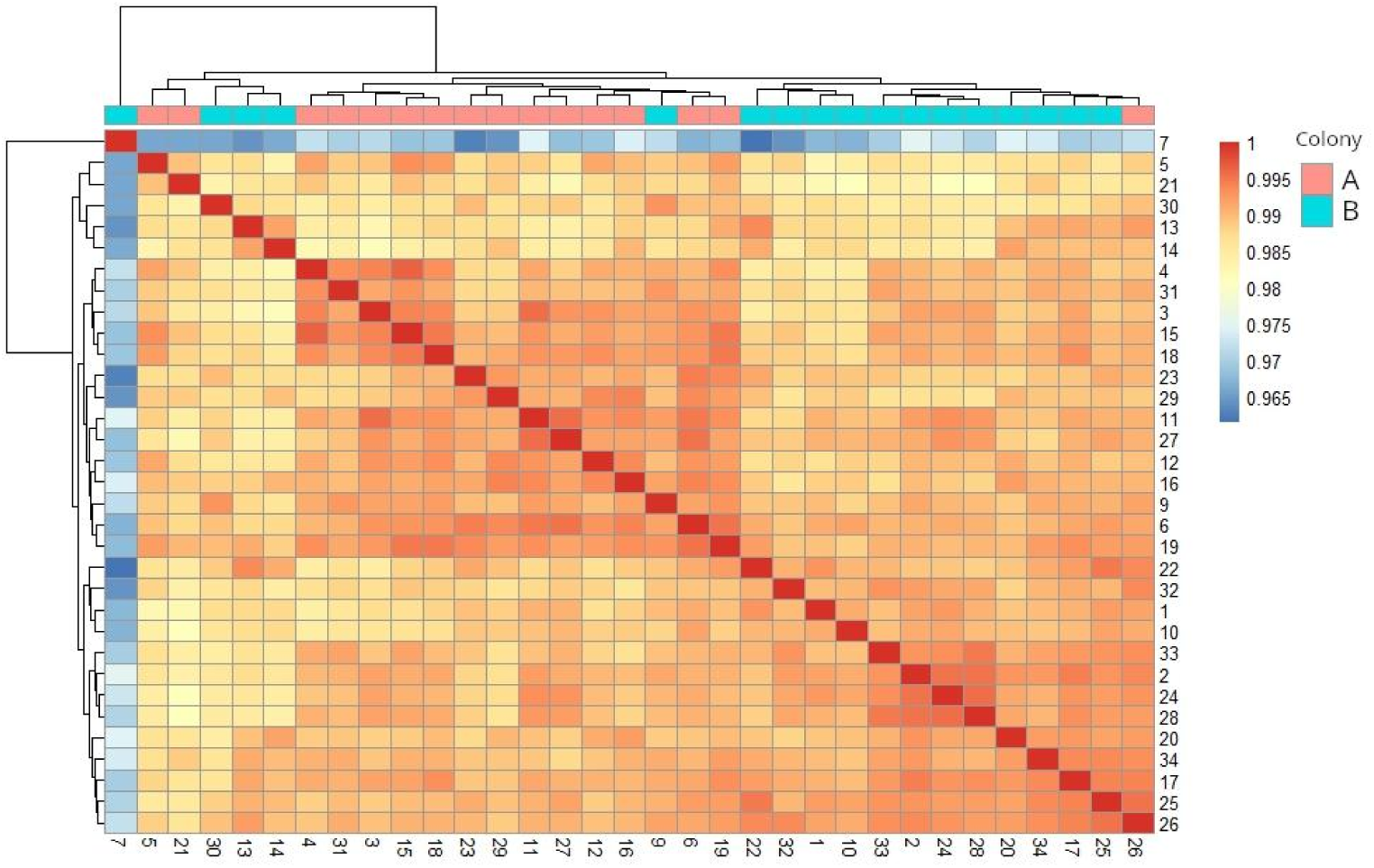
Heatmap showing hierarchical clustering of mushroom body gene profiles of the 12,332 genes from the 33 honey bee samples.

Given these observations, we chose to assess the transcriptomic data for Colony A separately, allowing us to control for confounding genetic and environmental factors. Specifically, the transcriptomic profile of the mushroom bodies changed as a result of high vs. low viral loads while disregarding learning performance. This analysis revealed a set of 50 genes that were significantly differentially expressed (Benjamini-Hochberg-adjusted *p* < 0.05). Of these, 13 genes were expressed with an absolute log_2_ fold change of one or more (|log₂FC| > 1) (Figure 4; Table 4). GO enrichment analysis revealed that key biological processes were overrepresented among these genes (Bonferroni-adjusted *p* < 0.001): GO:0042742/defense response to bacterium, GO:0002376/immune system process, and GO:0045087/innate immune response. The same three genes were significant in each group: *Apid1*, LOC406142, and LOC406144. KEGG pathway analysis identified enrichment in pathways associated with phototransduction and neuroactive ligand-receptor interaction, but neither of these results was significant following the Bonferroni correction (see Table S2).

**Figure 4.**
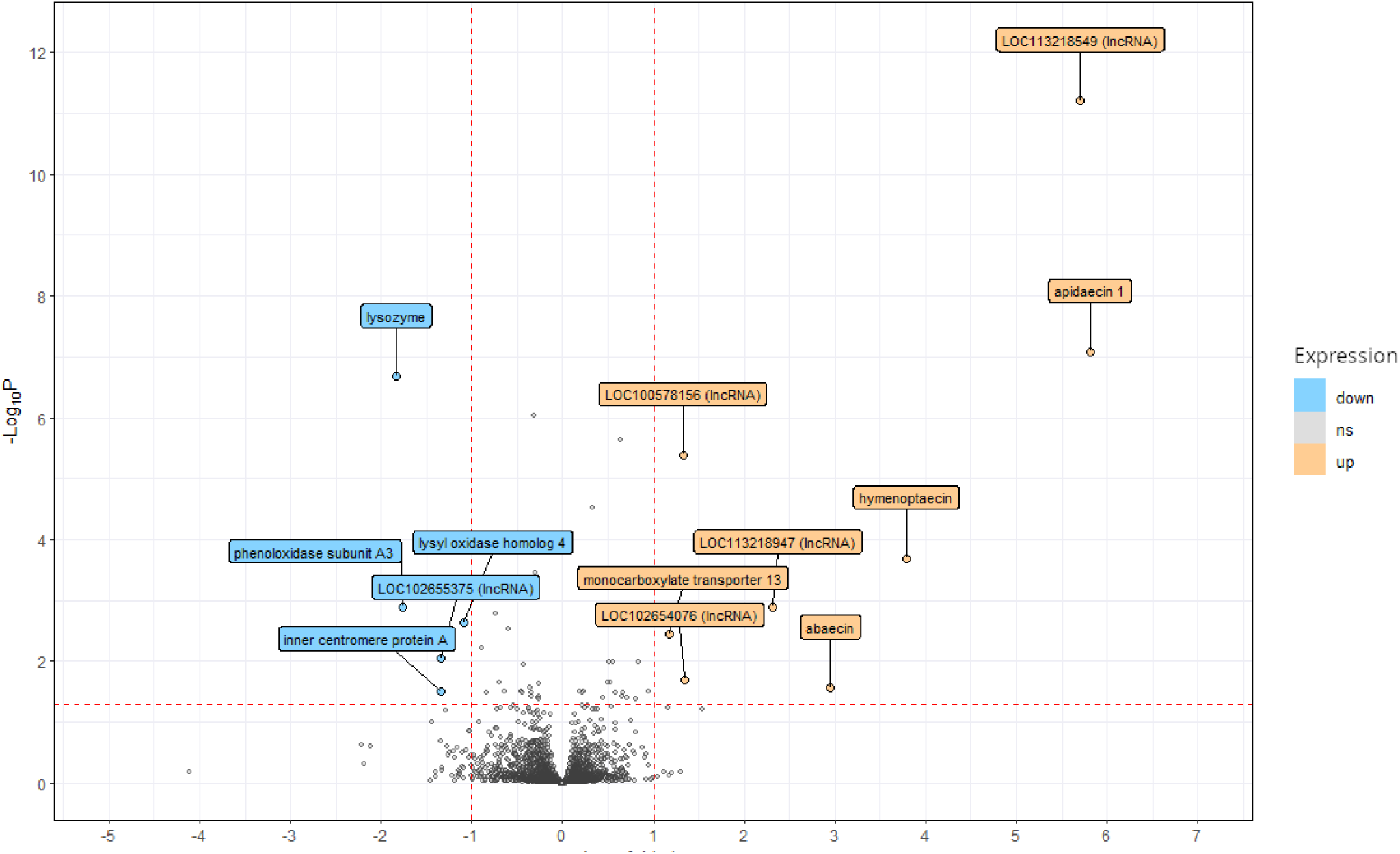
Volcano plot showing significantly expressed genes (Benjamini-Hochberg-adjusted *p* < 0.05, horizontal red dotted line) with an absolute log_2_ fold change of 1 or more (|log₂FC| > 1) (vertical red dotted lines) in bees from Colony A. Genes in orange are more highly expressed in bees with high viral loads (≥ log_10_ 5 DWV GE), while genes in blue are expressed at lower level in bees with high viral loads. Both measurements reported with reference to Colony A bees with low viral loads (≤ log_10_ 4 DWV GE).

**Table 4.**
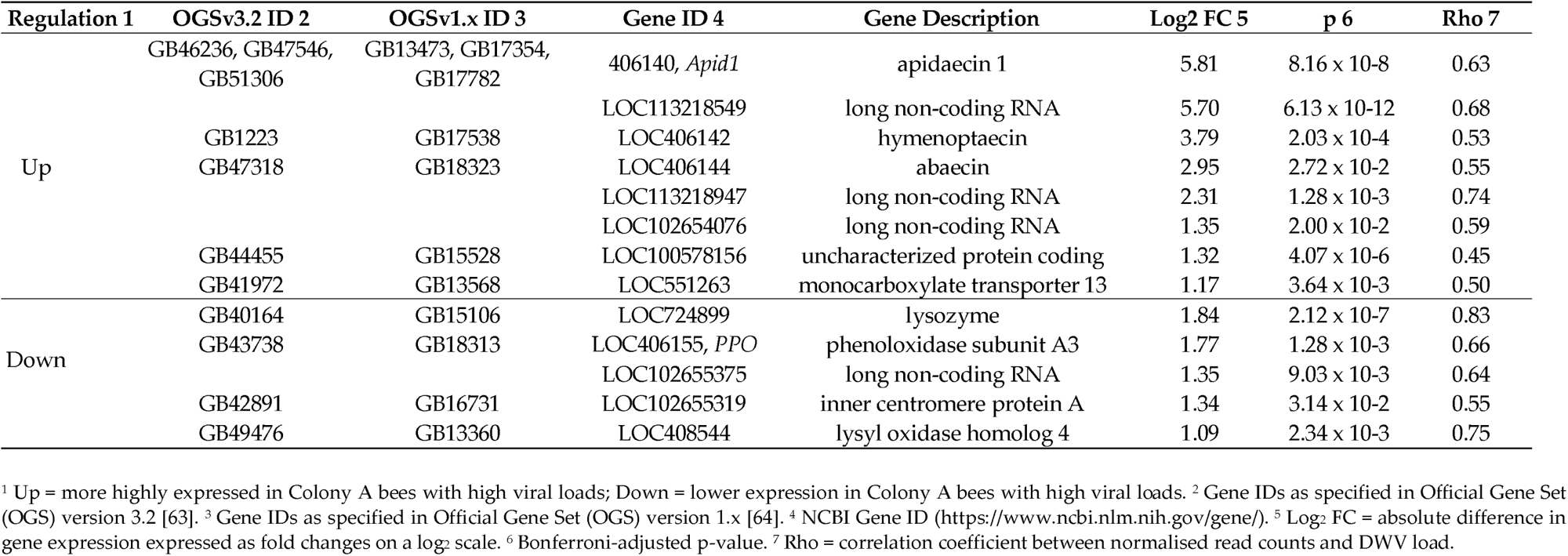
List of genes differentially expressed in bees from Colony A when comparing individuals with high vs. low viral loads (DWV ≥ log_10_ 5 and ≤ log_10_ 4, respectively)

### 3.4. Correlation between gene counts and viral loads

The analysis of the correlations (Spearman’s rank correlation coefficient) between gene counts and viral loads present in the mushroom bodies revealed some interesting patterns for both sets of data. Four genes showed strong correlation in the Poor Learners dataset (rho > 0.75, *p* < 0.001): LOC724642, LOC411159, LOC726505, and LOC107965291. Figure 5 shows plots for LOC724642/F-box/LRR-repeat protein 3 (panel a) and LOC107965291/a non-coding RNA (panel b). On the other hand, three genes showed strong correlation in the Colony A dataset (rho > 0.74, *p* < 0.001): LOC724899, LOC408544, and LOC113218947. Figure 5 shows plots for LOC724899/Lysozyme (panel c) and LOC113218947/ a non-coding RNA (panel d). Figures S2–S5 show correlation plots for allgenes listed in tables 3 and 4.

**Figure 5.**
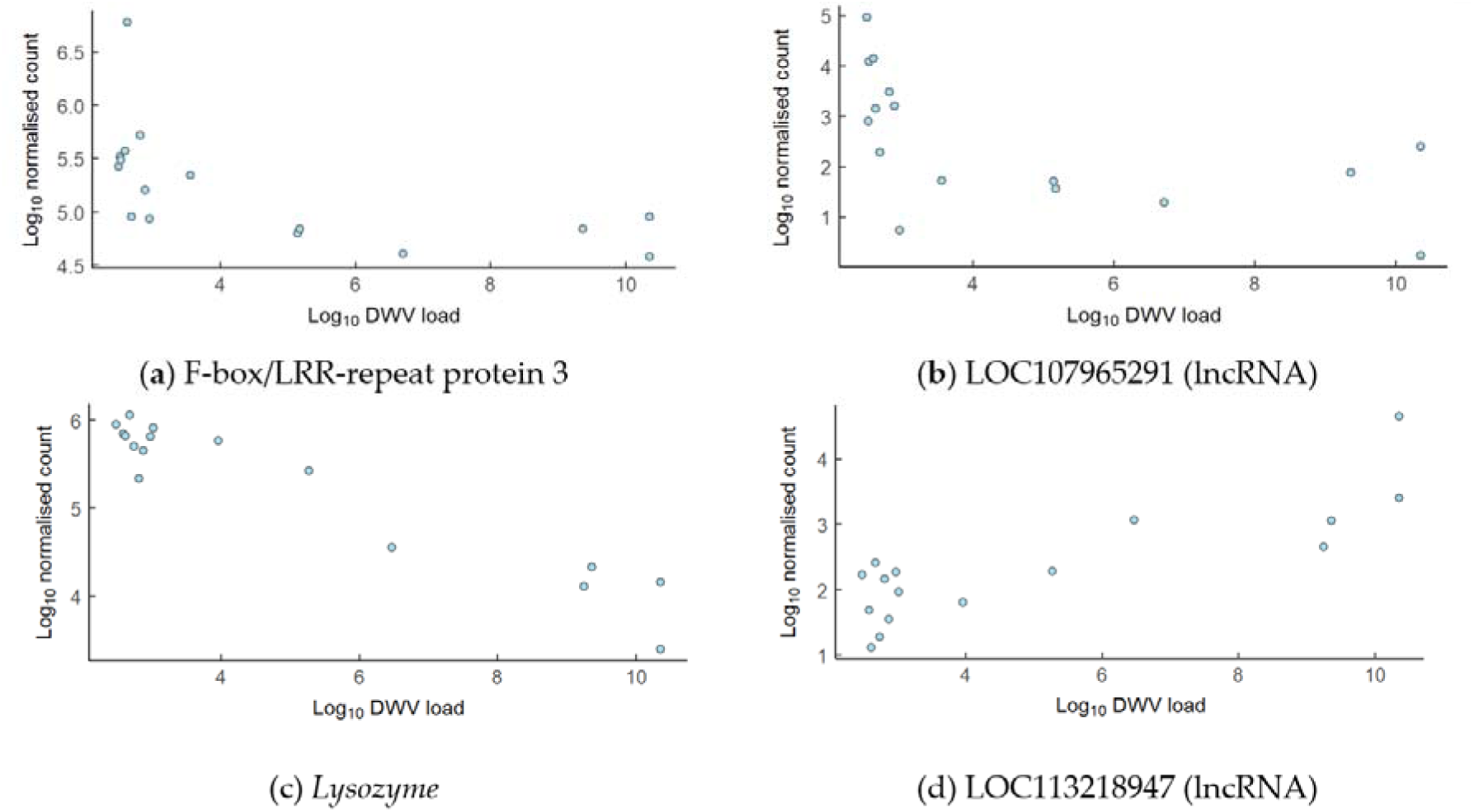
Highly significant correlations between gene counts and mushroom body viral loads for bees categorised as Poor Learners (a and b) and from Colony A (c and d). (a) LOC724642, F-box/LRR-repeat protein 3; Spearman’s Rank Correlation: rho = –0.79, *n* = 16, *p* < 0.001 (b) LOC107965291, a non-coding RNA; Spearman Rank correlation: rho = –0.75, *n* = 16, *p* < 0.001 (c) LOC274899, Lysozyme; Spearman Rank Correlation: rho = –0.83, *n* = 16, *p* < 0.001 (d) LOC113218947, a non-coding RNA; Spearman Rank correlation: rho = 0.74, *n* = 16, *p* < 0.001

### 3.5. Gene Regulatory Networks

GRN analysis of Colony A identified potential targets for lncRNAs LOC102654076, LOC102655375, LOC113218549, and LOC113218947; and analysis of the Poor Learners identified potential targets for lncRNAs LOC113218549, *and* LOC107965291. Table 5 lists the proposed target of lncRNA LOC113218549, which is common to both outputs; the other lncRNAs and their targets are shown in Tables S2-S5.

**Table 5.**
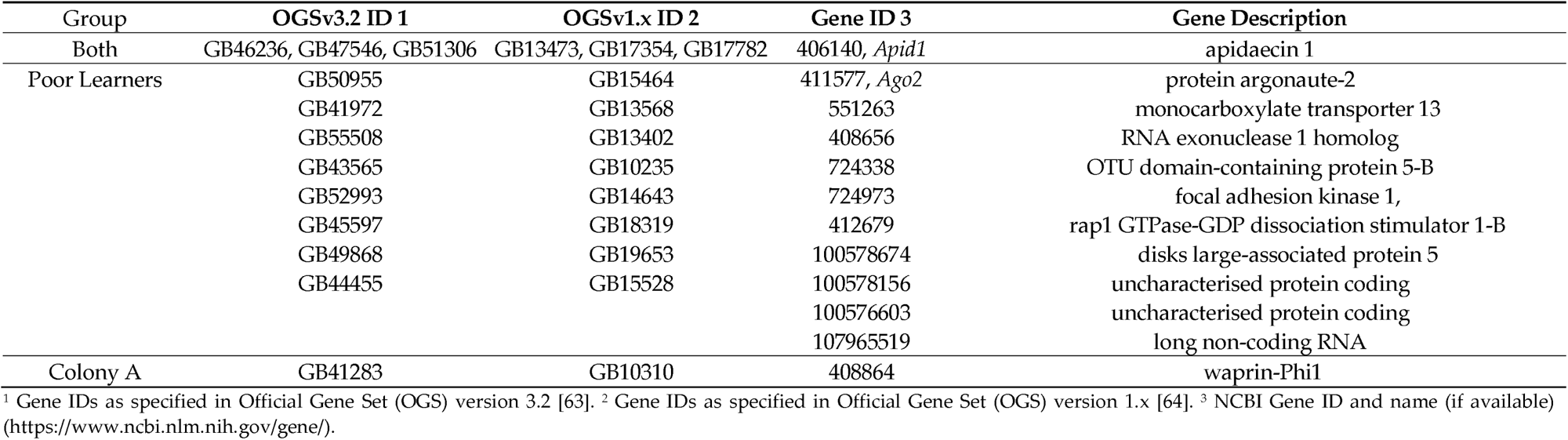
Genes identified as potential targets of lncRNA LOC113218549.

## 4. Discussion

A wealth of honey bee studies have examined the impact of viral infection on abdomen or whole body transcriptomes, and analysis of brain transcriptomes has become more common in recent years. *Pizzorno* et al. [38] have recently specifically explored DEGs in isolated brain tissue from honey bees with artificially-induced DWV infections, and in this study we have built on these findings by investigating the impact of naturally-occurring DWV infections on gene expression in honey bee foragers within a particular region of the brain—the mushroom bodies. Our initial analyses revealed that global gene expression patterns were primarily influenced by colony of origin, with Colony A showing distinct clustering patterns, higher DWV prevalence, and elevated viral loads. This led us to analyse bees from Colony A and Colony B separately to remove potential colony Background effects that could influence our results. The Colony A analysis revealed a distinct set of differentially expressed genes associated with higher DWV loads, including several immune-related genes and long non-coding RNAs (lncRNAs). Following these initial analyses, more-nuanced comparisons uncovered significant transcriptomic changes associated with higher DWV loads, particularly in bees exhibiting poor learning performance. In Poor Learners, high viral loads were linked to the differential expression of genes involved in immune response and sensory processing, including upregulation of antimicrobial peptides such as *apidaecin* and *hymenoptaecin*, and downregulation of genes related to neural signaling and metabolism. In contrast, no significant gene expression changes were observed in Good Learners when comparing high and low viral loads. Additionally, across both Poor Learners and Colony A bees, we found strong correlations between DWV load and the expression of specific genes, including lncRNAs. We used these lncRNAs to construct gene regulatory networks (GRNs) highlighting their potential regulatory role in modulating the honey bee brain’s response to viral infection.Notably, we observed no obvious transcriptomic signature in the mushroom bodies associated with learning performance, as revealed by our principal component analysis. This appears counterintuitive, given frequent reports linking learning performance with mushroom body activity and gene expression. [65–68]. However, since this study was concerned with the link between natural DWV load and simple associative learning, the procedure may not have been the most suitable to detect these effects. For example, the most suitable timing of capture and dissection of bees involved in learning tasks will vary according to needs [69,70]. Rittschof [71] cautions that although patterns of expression may indicate a tendency towards certain behaviours, that does not translate to actual performance of these behaviours. It is also possible that for a simple cognitive task such as associative learning, insufficient demands were placed on the mushroom bodies in order to yield detectable gene expression levels. It is possible that other brain regions play a larger role in regulating the response to such a simple task—for example the optic lobes or antennal lobes, more directly linked to the specific cues (visual and olfactory, respectively) presented during a PER assay. Szymanski et al. [42], who also investigated learning performance of bees carrying natural DWV infections, did not see any change in gene expression with simple associative learning tasks, but did note changes with more complex reversal learning. Notably, Szymanski et al. [42] targeted a specific group of GABA-related genes with a RT-qPCR approach, while we assessed the whole transcriptome of the honey bee. Interestingly, there were no GABA-related genes in our lists of differentially expressed genes. This could be due to differences in the complexity of the cognitive assay that we adopted, as discussed above, or due to differences in sensitivity between the RT-qPCR assay and an RNA-seq approach.

The most interesting discovery from our study was the detection of a clear transcriptomic signature associated with DWV infection in the mushroom bodies, evident in both the Poor Learner and Colony A datasets. DWV is known to actively replicate within the brain, including within the mushroom bodies [35], and we show here that immune genes are significantly expressed in this region of the honey bee brain. In both datasets, we observed upregulation of the antimicrobial peptides (AMPs) *apidaecin* and *hymenoptaecin*, while *abaecin* was upregulated in the Colony A dataset alone, suggesting possible variation in immune gene activation according to genetic and/or environmental background. Consistent with these findings, GO analysis revealed significant overrepresentation of immune-related categories in both groups, supporting the proposition that immune activation takes place within the mushroom bodies and that the insect brain therefore is not such an “immune-privileged” organ as initially thought [72–74]. In contrast to the overexpression of AMPs, another important immune-related gene, *lysozyme* (LOC724899), was down-regulated in bees carrying high viral loads from the Colony A dataset, showing a strong inverse relationship with viral load in our correlation analyses. Lysozyme’s role as an antibacterial, anti-viral, and immunomodulatory agent is well-established in both vertebrates and invertebrates [72,75–77]. Our findings align with previous observations that DWV infection can suppress lysozyme-mediated immunity in honey bees [78,79], likely by interfering with NF-κB signalling [80]. This dampening of the immune response mirrors similar observations reported for both invertebrates [81,82] and vertebrates [83], where infections by viruses have been linked to suppression of immune functions. The inverse relationship between viral load and lysozyme expression suggests a tightly-linked mechanism by which DWV weakens the host’s innate defences. In addition, we detected another set of genes generally well-known for their role as immune-regulators in insects, that were differentially expressed in the mushroom bodies of bees in our study. LOC100578156, orthologue of *Drosophila pirk*, a negative regulator of the immune deficiency pathway [84], was upregulated in the Colony A group. Flies that overexpress *pirk* are more susceptible to gram-negative bacterial infection, and the same gene has previously been identified as an immune effector in *A. mellifera*, where it is over-expressed in bees with Black queen cell virus [85]. Phenoloxidase subunit A3 (*PPO*) was significantly downregulated in Colony A bees with high viral loads. PPO is a well-characterised and crucial component of the insect immune system [86]. Its expression is known to vary according to developmental stage, where it is typically upregulated in white-eyed pupae, downregulated in brown-eyed pupae, and shows varying regulation in adults [38,87–90]. Intriguingly, LOC100577198, an uncharacterised protein-coding gene which was downregulated in the Poor Learner group in the present study, has also previously been identified as an AMP involved in detoxification and immunity [91].

In the Poor Learners dataset, we noted several downregulated genes associated with neural functions in insects and other invertebrates: Crustacean cardioactive peptide receptor, a highly conserved insect neuropeptide responsible for varied biological functions in a range of species [92,93]; Homeotic Protein Deformed, a Hox transcription factor that is crucial during embryonic development [94,95], which has been observed upregulated in response to insecticide exposure [96]; Krüppel-like factor 10, a transcriptional regulator which has been implicated in immune modulation [97,98]; and *Hbg3*, which encodes the alpha-glucosidase enzyme, a component of carbohydrate metabolism and whose expression is associated with the transition from nursing to foraging. Interestingly, the upregulation of *Hbg3* has also been noted in precocious foragers [31,99,100], and while other studies have shown both its upregulation [101,102] and downregulation [103,104] in response to insecticides and pathogens, its downregulation here in affected bees may demonstrate how viral infection can impact foraging.

Although the present study did not seek to identify lncRNAs, several were noted in both the Poor Learner and Colony A datasets. The expression of two of these lncRNAs showed strong correlation with viral load, further demonstrating their importance in response to DWV infection. It is well established that lncRNAs are associated with a range of biological functions, including regulation [105], and the results of our GRN analysis are intriguing, particularly the presence of *apidaecin* and *argonaute-2* as potential targets of LOC113218549. *Argonaute-2* is a key component of the short-interfering RNA (siRNA) pathway, involved in antiviral defence in invertebrates [106], and its upregulation has been observed in response to DWV infection [38,107–109] and other viruses [106,110]. The observation that in both datasets, the lncRNA LOC113218549 is the most strongly significantly differentially expressed gene after *apidaecin* is compelling.

Some limitations of this study should be recognised. First, our analysis was limited to two colonies, and as such caution should be exercised when generalising our findings to colonies existing in a variety of genetic and ecological contexts. Future studies could incorporate a wider range of settings in order to further elucidate the impact of these factors. Second, we examined the bees’ responses to PER, a relatively simple associative learning task. More refined or advanced challenges may reveal subtle or nuanced impacts of DWV on cognition that this study has failed to uncover. Third, we have considered a single time point post-infection, which cannot capture the probable varying temporal transcriptomic responses to DWV infection. Finally, our discovery of lncRNAs is intriguing, but further work will be required in order to validate the functional roles of these.

## 5. Conclusions

In this study we explored the neurotranscriptomic responses of honey bee mushroom bodies under naturally occurring DWV infections. We noted a distinct effect of colony of origin on differential gene expression: this clear separation between Colony A and Colony B samples is likely due to a combination of varying factors, such as quality of forage, Varroa presence/abundance and DWV loads, and genetic background. Additionally, Colony B received miticide treatments, which may have affected the mode of transmission in the colonies—by Varroa mites in Colony A and oral transmission in Colony B. We also noted a greater contrast in gene expression in bees with impaired associative learning abilities when examining differences in viral loads, despite no clear correlation between cognitive performance and transcriptomic signatures. These varying results demonstrate the importance of considering a combination of pathogen load, transmission route, treatments, neurotranscriptomic responses, and behaviour in order to understand the impact of viral pathogens on cognition.

Our work highlights the benefits of analyses carried out on bees with naturally-occurring DWV infections over the induction of infections either with abdominal injections or by feeding. A comparison of the effects of these two routes of infection, and confirmation of which more closely mimics natural infections, would be informative, since it would facilitate future studies that wish to examine the effects of natural infections. Moreover, natural infections such as those in our study occur during the larval or pupal stages, allowing potential effects on the developing brain to be assessed, unlike studies involving DWV injection into adult bees, whose brains are already fully developed. We have also emphasised how our approach here has further refined studies of infection and gene expression in the honey bee brain, by focussing on the mushroom bodies. A further refinement would be to use single-cell sequencing to compare infected and non-infected neurons within the mushroom bodies.

## Supplementary Materials

The following supporting information can be downloaded at: www.mdpi.com/xxx/, Figure S1: Scree plot showing the variance explained by each principal component, Figure S2: Correlations between gene counts of upregulated genes and mushroom body viral loads for bees categorised as Poor Learners, Figure S3: Correlations between gene of downregulated genes and mushroom body viral loads for bees categorised as Poor Learners, Figure S4: Correlations between gene counts of upregulated genes and mushroom body viral loads for bees from Colony A, Figure S5: Correlations between gene counts of downregulated genes and mushroom body viral loads for bees from Colony A, Table S1: KEGG pathways identified via overrepresentation analysis of Poor Learners, Table S2: KEGG pathways identified via overrepresentation analysis of Colony A bees, Table S3: Genes identified as potential targets of lncRNAs

## Author Contributions

Conceptualization, S.L., A.B. and F.M.; methodology, S.L. and L.D.; software, S.L.; formal analysis, S.L.; investigation, S.L. and L.D.; resources, A.B. and F.M.; data curation, S.L.; writing—original draft preparation, S.L. and F.M.; writing—review and editing, S.L., L.D., A.B. and F.M.; visualization, S.L.; supervision, F.M.; project administration, F.M.; funding acquisition, A.B. and F.M. All authors have read and agreed to the published version of the manuscript.

## Data Availability Statement

The original data presented in the study will be openly available in the NCBI Sequence Read Archive (SRA) at https://www.ncbi.nlm.nih.gov/sra/PRJNA1261397. The code used for analysis will be made available via GitHub at https://github.com/simonedwardloughran/neurotranscriptomics-dwv

## Acknowledgments

We would like to thank Ewan Campbell for beekeeping support and the CGBEM (Centre for Genome Enabled Biology and Medicine) at the University of Aberdeen for sequencing and processing of raw data.

## Disclaimer/Publisher’s Note

The statements, opinions and data contained in all publications are solely those of the individual author(s) and contributor(s) and not of MDPI and/or the editor(s). MDPI and/or the editor(s) disclaim responsibility for any injury to people or property resulting from any ideas, methods, instructions or products referred to in the content.

## References

1. Schippers M-P, Dukas R, McClelland GB. Lifetime- and caste-specific changes in flight metabolic rate and muscle biochemistry of honeybees, Apis mellifera. J Comp Physiol B. 2010;180: 45–55.

2. Rothe U, Nachtigall W. Flight of the honey bee: IV. Respiratory quotients and metabolic rates during sitting, walking and flying. J Comp Physiol B. 1989;158: 739–749.

3. Doussot C, Purdy J, Lihoreau M. Navigation: Cognition, learning, and memory. The Foraging Behavior of the Honey Bee (Apis mellifera, L). Elsevier; 2024. pp. 85–104.

4. Bullinger E, Greggers U, Menzel R. Generalization of navigation memory in honeybees. Front Behav Neurosci. 2023;17: 1070957.

5. Klein S, Cabirol A, Devaud J-M, Barron AB, Lihoreau M. Why Bees Are So Vulnerable to Environmental Stressors. Trends Ecol Evol. 2017;32: 268–278.

6. Bordier C, Suchail S, Pioz M, Devaud JM, Collet C, Charreton M, et al. Stress response in honeybees is associated with changes in task-related physiology and energetic metabolism. J Insect Physiol. 2017;98: 47–54.

7. Koch H, Brown MJ, Stevenson PC. The role of disease in bee foraging ecology. Curr Opin Insect Sci. 2017;21: 60–67.

8. Gómez-Moracho T, Heeb P, Lihoreau M. Effects of parasites and pathogens on bee cognition: Bee parasites, pathogens and cognition. Ecol Entomol. 2017;42: 51–64.

9. Grab H, Branstetter MG, Amon N, Urban-Mead KR, Park MG, Gibbs J, et al. Agriculturally dominated landscapes reduce bee phylogenetic diversity and pollination services. Science. 2019;363: 282–284.

10. Kammerer M, Goslee SC, Douglas MR, Tooker JF, Grozinger CM. Wild bees as winners and losers: Relative impacts of landscape composition, quality, and climate. Glob Chang Biol. 2021;27: 1250–1265.

11. Siviter H, Koricheva J, Brown MJF, Leadbeater E. Quantifying the impact of pesticides on learning and memory in bees. J Appl Ecol. 2018;55: 2812–2821.

12. Belzunces LP, Tchamitchian S, Brunet J-L. Neural effects of insecticides in the honey bee. Apidologie (Celle). 2012;43: 348–370.

13. Peng Y-C, Yang E-C. Sublethal dosage of imidacloprid reduces the microglomerular density of honey bee mushroom bodies. Sci Rep. 2016;6: 19298.

14. Palmer MJ, Moffat C, Saranzewa N, Harvey J, Wright GA, Connolly CN. Cholinergic pesticides cause mushroom body neuronal inactivation in honeybees. Nat Commun. 2013;4: 1634.

15. French SK, Pepinelli M, Conflitti IM, Jamieson A, Higo H, Common J, et al. Honey bee stressor networks are complex and dependent on crop and region. Curr Biol. 2024. doi:10.1016/j.cub.2024.03.039

16. Straub L, Minnameyer A, Strobl V, Kolari E, Friedli A, Kalbermatten I, et al. From antagonism to synergism: Extreme differences in stressor interactions in one species. Sci Rep. 2020;10: 4667.

17. El-Seedi HR, Ahmed HR, El-Wahed AAA, Saeed A, Algethami AF, Attia NF, et al. Bee stressors from an immunological perspective and strategies to improve bee health. Vet Sci. 2022;9: 199.

18. Even N, Devaud J-M, Barron AB. General stress responses in the honey bee. Insects. 2012;3: 1271–1298.

19. de Miranda JR, Bailey L, Ball BV, Blanchard P, Budge GE, Chejanovsky N, et al. Standard methods for virus research inApis mellifera. J Apic Res. 2013;52: 1–56.

20. Beaurepaire A, Piot N, Doublet V, Antunez K, Campbell E, Chantawannakul P, et al. Diversity and global distribution of viruses of the western honey bee, Apis mellifera. Insects. 2020;11: 239.

21. McMenamin AJ, Flenniken ML. Recently identified bee viruses and their impact on bee pollinators. Curr Opin Insect Sci. 2018;26: 120–129.

22. Brutscher LM, McMenamin AJ, Flenniken ML. The buzz about honey bee viruses. PLoS Pathog. 2016;12: e1005757.

23. Wilfert L, Long G, Leggett HC, Schmid-Hempel P, Butlin R, Martin SJM, et al. Deformed wing virus is a recent global epidemic in honeybees driven by Varroa mites. Science. 2016;351: 594–597.

24. Kevill JL, de Souza FS, Sharples C, Oliver R, Schroeder DC, Martin SJ. DWV-A lethal to honey bees (Apis mellifera): A colony level survey of DWV variants (A, B, and C) in England, Wales, and 32 states across the US. Viruses. 2019;11: 426.

25. Ramsey SD, Ochoa R, Bauchan G, Gulbronson C, Mowery JD, Cohen A, et al. Varroa destructor feeds primarily on honey bee fat body tissue and not hemolymph. Proc Natl Acad Sci U S 2019;116: 1792–1801.

26. Martin SJ, Highfield AC, Brettell L, Villalobos EM, Budge GE, Powell M, et al. Global honey bee viral landscape altered by a parasitic mite. Science. 2012;336: 1304–1306.

27. Santillán-Galicia MT, Ball BV, Clark SJ, Alderson PG. Transmission of deformed wing virus and slow paralysis virus to adult bees (Apis melliferaL.) byVarroa destructor. J Apic Res. 2010;49: 141–148.

28. Koziy RV, Wood SC, Kozii IV, van Rensburg CJ, Moshynskyy I, Dvylyuk I, et al. Deformed wing virus infection in honey bees (Apis mellifera L.). Vet Pathol. 2019;56: 636–641.

29. de Miranda JR, Genersch E. Deformed wing virus. J Invertebr Pathol. 2010;103 Suppl 1: S48–61.

30. de Miranda JR, Fries I. Venereal and vertical transmission of deformed wing virus in honeybees (Apis mellifera L.). J Invertebr Pathol. 2008;98: 184–189.

31. Traniello IM, Bukhari SA, Kevill J, Ahmed AC, Hamilton AR, Naeger NL, et al. Meta-analysis of honey bee neurogenomic response links Deformed wing virus type A to precocious behavioral maturation. Sci Rep. 2020;10: 3101.

32. Wells T, Wolf S, Nicholls E, Groll H, Lim KS, Clark SJ, et al. Flight performance of actively foraging honey bees is reduced by a common pathogen. Environ Microbiol Rep. 2016;8: 728–737.

33. Benaets K, Van Geystelen A, Cardoen D, De Smet L, de Graaf DC, Schoofs L, et al. Covert deformed wing virus infections have long-term deleterious effects on honeybee foraging and survival. Proc Biol Sci. 2017;284: 20162149.

34. Penn HJ, Simone-Finstrom MD, de Guzman LI, Tokarz PG, Dickens R. Colony-level viral load influences collective foraging in honey bees. Front Insect Sci. 2022;2: 894482.

35. Shah KS, Evans EC, Pizzorno MC. Localization of deformed wing virus (DWV) in the brains of the honeybee, Apis mellifera Linnaeus. Virol J. 2009;6: 182.

36. Doublet V, Poeschl Y, Gogol-Döring A, Alaux C, Annoscia D, Aurori C, et al. Unity in defence: honeybee workers exhibit conserved molecular responses to diverse pathogens. BMC Genomics. 2017;18: 207.

37. Zhao X, Liu Y. Current knowledge on bee innate immunity based on genomics and transcriptomics. Int J Mol Sci. 2022;23: 14278.

38. Pizzorno MC, Field K, Kobokovich AL, Martin PL, Gupta RA, Mammone R, et al. Transcriptomic Responses of the Honey Bee Brain to Infection with Deformed Wing Virus. Viruses. 2021;13. doi:10.3390/v13020287

39. Iqbal J, Mueller U. Virus infection causes specific learning deficits in honeybee foragers. Proc Biol Sci. 2007;274: 1517–1521.

40. Chen P, Lu Y-H, Lin Y-H, Wu C-P, Tang C-K, Wei S-C, et al. Deformed wing virus infection affects the neurological function of Apis mellifera by altering extracellular adenosine signaling. Insect Biochem Mol Biol. 2021;139: 103674.

41. Farris SM, Robinson GE, Davis RL, Fahrbach SE. Larval and pupal development of the mushroom bodies in the honey bee, Apis mellifera. J Comp Neurol. 1999;414: 97–113.

42. Szymański S, Baracchi D, Dingle L, Bowman AS, Manfredini F. Learning performance and GABAergic pathway link to deformed wing virus in the mushroom bodies of naturally infected honey bees. J Exp Biol. 2024;227: jeb246766.

43. Zanni V, Frizzera D, Marroni F, Seffin E, Annoscia D, Nazzi F. Age-related response to mite parasitization and viral infection in the honey bee suggests a trade-off between growth and immunity. PLoS One. 2023;18: e0288821.

44. Woodford L, Christie CR, Campbell EM, Budge GE, Bowman AS, Evans DJ. Quantitative and qualitative changes in the Deformed wing virus population in honey bees associated with the introduction or removal of Varroa destructor. Viruses. 2022;14: 1597.

45. Mota T, Giurfa M. Multiple reversal olfactory learning in honeybees. Front Behav Neurosci. 2010;4: 1896.

46. Bradford EL, Christie CR, Campbell EM, Bowman AS. A real-time PCR method for quantification of the total and major variant strains of the deformed wing virus. PLoS One. 2017;12: e0190017.

47. Krueger F. TrimGalore: A wrapper around Cutadapt and FastQC to consistently apply adapter and quality trimming to FastQ files, with extra functionality for RRBS data. Github; 2019. Available: https://github.com/FelixKrueger/TrimGalore

48. Andrews. Babraham Bioinformatics - FastQC A Quality Control tool for High Throughput Sequence Data. 2010 [cited 26 Jan 2023]. Available: https://www.bioinformatics.babraham.ac.uk/projects/fastqc/

49. Kim D, Langmead B, Salzberg SL. HISAT: a fast spliced aligner with low memory requirements. Nat Methods. 2015;12: 357–360.

50. Norton AM, Remnant EJ, Buchmann G, Beekman M. Accumulation and Competition Amongst Deformed Wing Virus Genotypes in Naïve Australian Honeybees Provides Insight Into the Increasing Global Prevalence of Genotype B. Front Microbiol. 2020;11: 620.

51. Dalmon A, Gayral P, Decante D, Klopp C, Bigot D, Thomasson M, et al. Viruses in the Invasive Hornet Vespa velutina. Viruses. 2019;11. doi:10.3390/v11111041

52. Liao Y, Smyth GK, Shi W. featureCounts: an efficient general purpose program for assigning sequence reads to genomic features. Bioinformatics. 2014;30: 923–930.

53. Soneson C, Robinson MD. Bias, robustness and scalability in single-cell differential expression analysis. Nat Methods. 2018;15: 255–261.

54. Satija R, Farrell JA, Gennert D, Schier AF, Regev A. Spatial reconstruction of single-cell gene expression data. Nat Biotechnol. 2015;33: 495–502.

55. Love MI, Huber W, Anders S. Moderated estimation of fold change and dispersion for RNA-seq data with DESeq2. Genome Biol. 2014;15: 550.

56. Anders S, Huber W. Differential expression analysis for sequence count data. Genome Biol. 2010;11: R106.

57. Rousseeuw PJ. Silhouettes: A graphical aid to the interpretation and validation of cluster analysis. J Comput Appl Math. 1987;20: 53–65.

58. Robinson MD, McCarthy DJ, Smyth GK. edgeR: a Bioconductor package for differential expression analysis of digital gene expression data. Bioinformatics. 2010;26: 139–140.

59. Geistlinger L, Csaba G, Zimmer R. Bioconductor’s EnrichmentBrowser: seamless navigation through combined results of set- & network-based enrichment analysis. BMC Bioinformatics. 2016;17: 45.

60. Goeman JJ, Bühlmann P. Analyzing gene expression data in terms of gene sets: methodological issues. Bioinformatics. 2007;23: 980–987.

61. Ashburner M, Ball CA, Blake JA, Botstein D, Butler H, Cherry JM, et al. Gene ontology: tool for the unification of biology. The Gene Ontology Consortium. Nat Genet. 2000;25: 25–29.

62. Kanehisa M, Araki M, Goto S, Hattori M, Hirakawa M, Itoh M, et al. KEGG for linking genomes to life and the environment. Nucleic Acids Res. 2008;36: D480–4.

63. Elsik CG, Worley KC, Bennett AK, Beye M, Camara F, Childers CP, et al. Finding the missing honey bee genes: lessons learned from a genome upgrade. BMC Genomics. 2014;15: 86.

64. Honeybee Genome Sequencing Consortium. Insights into social insects from the genome of the honeybee Apis mellifera. Nature. 2006;443: 931–949.

65. Szyszka P, Galkin A, Menzel R. Associative and non-associative plasticity in kenyon cells of the honeybee mushroom body. Front Syst Neurosci. 2008;2: 3.

66. Giurfa M. Cognitive neuroethology: dissecting non-elemental learning in a honeybee brain. Curr Opin Neurobiol. 2003;13: 726–735.

67. Strube-Bloss MF, Nawrot MP, Menzel R. Mushroom body output neurons encode odor-reward associations. J Neurosci. 2011;31: 3129–3140.

68. Fahad Raza M, Anwar M, Husain A, Rizwan M, Li Z, Nie H, et al. Differential gene expression analysis following olfactory learning in honeybee (Apis mellifera L.). PLoS One. 2022;17: e0262441.

69. Rittschof CC, Hughes KA. Advancing behavioural genomics by considering timescale. Nat Commun. 2018;9: 489.

70. Clayton DF. The genomic action potential. Neurobiol Learn Mem. 2000;74: 185–216.

71. Rittschof CC. Sequential social experiences interact to modulate aggression but not brain gene expression in the honey bee (Apis mellifera). Front Zool. 2017;14: 16.

72. Feng M, Fei S, Zou J, Xia J, Lai W, Huang Y, et al. Single-nucleus sequencing of silkworm larval brain reveals the key role of lysozyme in the antiviral immune response in brain hemocytes. J Innate Immun. 2024;16: 173–187.

73. Contreras EG, Klämbt C. The Drosophila blood-brain barrier emerges as a model for understanding human brain diseases. Neurobiol Dis. 2023;180: 106071.

74. Lye SH, Chtarbanova S. Drosophila as a model to study brain innate immunity in health and disease. Int J Mol Sci. 2018;19: 3922.

75. Liu L, Jia X, Zhao X, Li T, Luo Z, Deng R, et al. In vitro PCR verification that lysozyme inhibits nucleic acid replication and transcription. Sci Rep. 2023;13: 6383.

76. Ragland SA, Criss AK. From bacterial killing to immune modulation: Recent insights into the functions of lysozyme. PLoS Pathog. 2017;13: e1006512.

77. Chen T-T, Tan L-R, Hu N, Dong Z-Q, Hu Z-G, Jiang Y-M, et al. C-lysozyme contributes to antiviral immunity in Bombyx mori against nucleopolyhedrovirus infection. J Insect Physiol. 2018;108: 54–60.

78. Ray AM, Tehel A, Rasgon JL, Paxton RJ, Grozinger CM. The intensity of the transcriptional response varies across infection with distinct viral strains in an insect host. BMC Genomics. 2025;26: 175.

79. Yang X, Cox-Foster DL. Impact of an ectoparasite on the immunity and pathology of an invertebrate: evidence for host immunosuppression and viral amplification. Proc Natl Acad Sci U S 2005;102: 7470–7475.

80. Di Prisco G, Annoscia D, Margiotta M, Ferrara R, Varricchio P, Zanni V, et al. A mutualistic symbiosis between a parasitic mite and a pathogenic virus undermines honey bee immunity and health. Proc Natl Acad Sci U S A. 2016;113: 3203–3208.

81. Palmer WH, Joosten J, Overheul GJ, Jansen PW, Vermeulen M, Obbard DJ, et al. Induction and suppression of NF-κB signalling by a DNA virus of Drosophila. J Virol. 2019;93: e01443–18.

82. Shelby KS, Cui L, Webb BA. Polydnavirus-mediated inhibition of lysozyme gene expression and the antibacterial response. Insect Mol Biol. 1998;7: 265–272.

83. Pang G, Clancy R, Cong M, Ortega M, Zhigang R, Reeves G. Influenza virus inhibits lysozyme secretion by sputum neutrophils in subjects with chronic bronchial sepsis. Am J Respir Crit Care Med. 2000;161: 718–722.

84. Kleino A, Myllymäki H, Kallio J, Vanha-aho L-M, Oksanen K, Ulvila J, et al. Pirk is a negative regulator of the Drosophila Imd pathway. J Immunol. 2008;180: 5413–5422.

85. Doublet V, Paxton RJ, McDonnell CM, Dubois E, Nidelet S, Moritz RFA, et al. Brain transcriptomes of honey bees (Apis mellifera) experimentally infected by two pathogens: Black queen cell virus and Nosema ceranae. Genom Data. 2016;10: 79–82.

86. González-Santoyo I, Córdoba-Aguilar A. Phenoloxidase: a key component of the insect immune system: Biochemical and evolutionary ecology of PO. Entomol Exp Appl. 2012;142: 1–16.

87. Tesovnik T, Zorc M, Gregorc A, Rinehart T, Adamczyk J, Narat M. Immune gene expression in developing honey bees (Apis mellifera L.) simultaneously exposed to imidacloprid and Varroa destructor in laboratory conditions. J Apic Res. 2019;58: 730–739.

88. Zaobidna EA, Żółtowska K, Łopieńska-Biernat E. Expression of the prophenoloxidase gene and phenoloxidase activity, during the development of Apis mellifera brood infected with Varroa destructor. J Apic Sci. 2015;59: 85–93.

89. Khongphinitbunjong K, de Guzman LI, Tarver MR, Rinderer TE, Chen Y, Chantawannakul P. Differential viral levels and immune gene expression in three stocks of Apis mellifera induced by different numbers of Varroa destructor. J Insect Physiol. 2015;72: 28–34.

90. Kuster RD, Boncristiani HF, Rueppell O. Immunogene and viral transcript dynamics during parasitic Varroa destructor mite infection of developing honey bee (Apis mellifera) pupae. J Exp Biol. 2014;217: 1710–1718.

91. Wu T, Gao J, Choi YS, Kim DW, Han B, Yang S, et al. Interaction of chlorothalonil and Varroa destructor on immature honey bees rearing in vitro. Sci Total Environ. 2023;904: 166302.

92. Zhou Y, Nagata S. Crustacean cardioactive peptide. Handbook of Hormones. Elsevier; 2021. pp. 795–797.

93. Stangier J, Hilbich C, Beyreuther K, Keller R. Unusual cardioactive peptide (CCAP) from pericardial organs of the shore crab Carcinus maenas. Proc Natl Acad Sci U S A. 1987;84: 575–579.

94. Abzhanov A, Kaufman TC. Homeotic genes and the arthropod head: expression patterns of the labial, proboscipedia, and Deformed genes in crustaceans and insects. Proc Natl Acad Sci U S A. 1999;96: 10224–10229.

95. Feng W, Li Y, Kratsios P. Emerging roles for Hox proteins in the last steps of neuronal development in worms, flies, and mice. Front Neurosci. 2021;15: 801791.

96. Fent K, Schmid M, Christen V. Global transcriptome analysis reveals relevant effects at environmental concentrations of cypermethrin in honey bees (Apis mellifera). Environ Pollut. 2020;259: 113715.

97. Kaczynski J, Cook T, Urrutia R. Sp1- and Krüppel-like transcription factors. Genome Biol. 2003;4: 206.

98. Kulkarni A, Pandey A, Trainor P, Carlisle S, Yu W, Kukutla P, et al. Aryl hydrocarbon receptor and Krüppel like factor 10 mediate a transcriptional axis modulating immune homeostasis in mosquitoes. Sci Rep. 2022;12: 6005.

99. Ueno T, Takeuchi H, Kawasaki K, Kubo T. Changes in the gene expression profiles of the hypopharyngeal gland of worker honeybees in association with worker behavior and hormonal factors. PLoS One. 2015;10: e0130206.

100. Colin T, Meikle WG, Wu X, Barron AB. Traces of a neonicotinoid induce precocious foraging and reduce foraging performance in honey bees. Environ Sci Technol. 2019;53: 8252–8261.

101. Dussaubat C, Brunet J-L, Higes M, Colbourne JK, Lopez J, Choi J-H, et al. Gut pathology and responses to the microsporidium Nosema ceranae in the honey bee Apis mellifera. PLoS One. 2012;7: e37017.

102. Fent K, Haltiner T, Kunz P, Christen V. Insecticides cause transcriptional alterations of endocrine related genes in the brain of honey bee foragers. Chemosphere. 2020;260: 127542.

103. Chen Y-R, Tzeng DTW, Yang E-C. Chronic effects of imidacloprid on honey bee worker development-molecular pathway perspectives. Int J Mol Sci. 2021;22: 11835.

104. de Castro Lippi IC, Lima YS, da Luz Scheffer J, Lunardi JS, Kadri SM, Alvarez MVN, et al. Transcriptomic analysis of the head reveals molecular mechanisms underlying topical imidacloprid effects on A. mellifera forager bees. Apidologie (Celle). 2025;56: 1–17.

105. Wang KC, Chang HY. Molecular mechanisms of long noncoding RNAs. Mol Cell. 2011;43: 904–914.

106. Brutscher LM, Flenniken ML. RNAi and antiviral defense in the honey bee. J Immunol Res. 2015;2015: 941897.

107. Zhao Y, Heerman M, Peng W, Evans JD, Rose R, DeGrandi-Hoffman G, et al. The dynamics of Deformed wing virus concentration and host defensive gene expression after Varroa mite parasitism in honey bees, Apis mellifera. Insects. 2019;10: 16.

108. Norton AM, Buchmann G, Ashe A, Watson OT, Beekman M, Remnant EJ. Deformed wing virus genotypes A and B do not elicit immunologically different responses in naïve honey bee hosts. Insect Mol Biol. 2025;34: 33–51.

109. Brutscher LM, Daughenbaugh KF, Flenniken ML. Virus and dsRNA-triggered transcriptional responses reveal key components of honey bee antiviral defense. Sci Rep. 2017;7: 6448.

110. Galbraith DA, Yang X, Niño EL, Yi S, Grozinger C. Parallel epigenomic and transcriptomic responses to viral infection in honey bees (Apis mellifera). PLoS Pathog. 2015;11: e1004713.

