## Supplementary Material for "Neurotranscriptomic profiling of DWV-infected honey bee foragers with different cognitive abilities"

**Supplementary Figure Legends:**

**Figure S1**: Scree plot showing the variance explained by each principal component

**Figure S2**: Correlations between gene counts of upregulated genes and mushroom body viral loads for bees categorised as Poor Learners. (a) LOC406140, *Apid1* (b) LOC113218549 (c) LOC406142, *hymenoptaecin*

**Figure S3**: Correlations between gene counts of downregulated genes and mushroom body viral loads for bees categorised as Poor Learners. (a) LOC102655185 (b) LOC102654257 (c) LOC107965291 (d) LOC406131, *Hbg3* (e) LOC724252, *Dfd* (f) LOC726505 (g) LOC410326 (h) LOC412829 (i) LOC726935 (j) LOC411159 (k) LOC724642 (l) LOC100577198

**Figure S4**: Correlations between gene counts of upregulated genes and mushroom body viral loads for bees from Colony A. (a) 406140, *Apid1* (b) LOC113218549 (c) LOC406142 (d) LOC406144 (e) LOC113218947 (f) LOC102654076 (g) LOC100578156 (h) LOC551263

**Figure S5**: Correlations between gene counts of downregulated genes and mushroom body viral loads for bees from Colony A. (a) LOC724899 (b) LOC406155, *PPO* (c) LOC102655375 (d) LOC102655319 (e) LOC408544

**Figure S1**

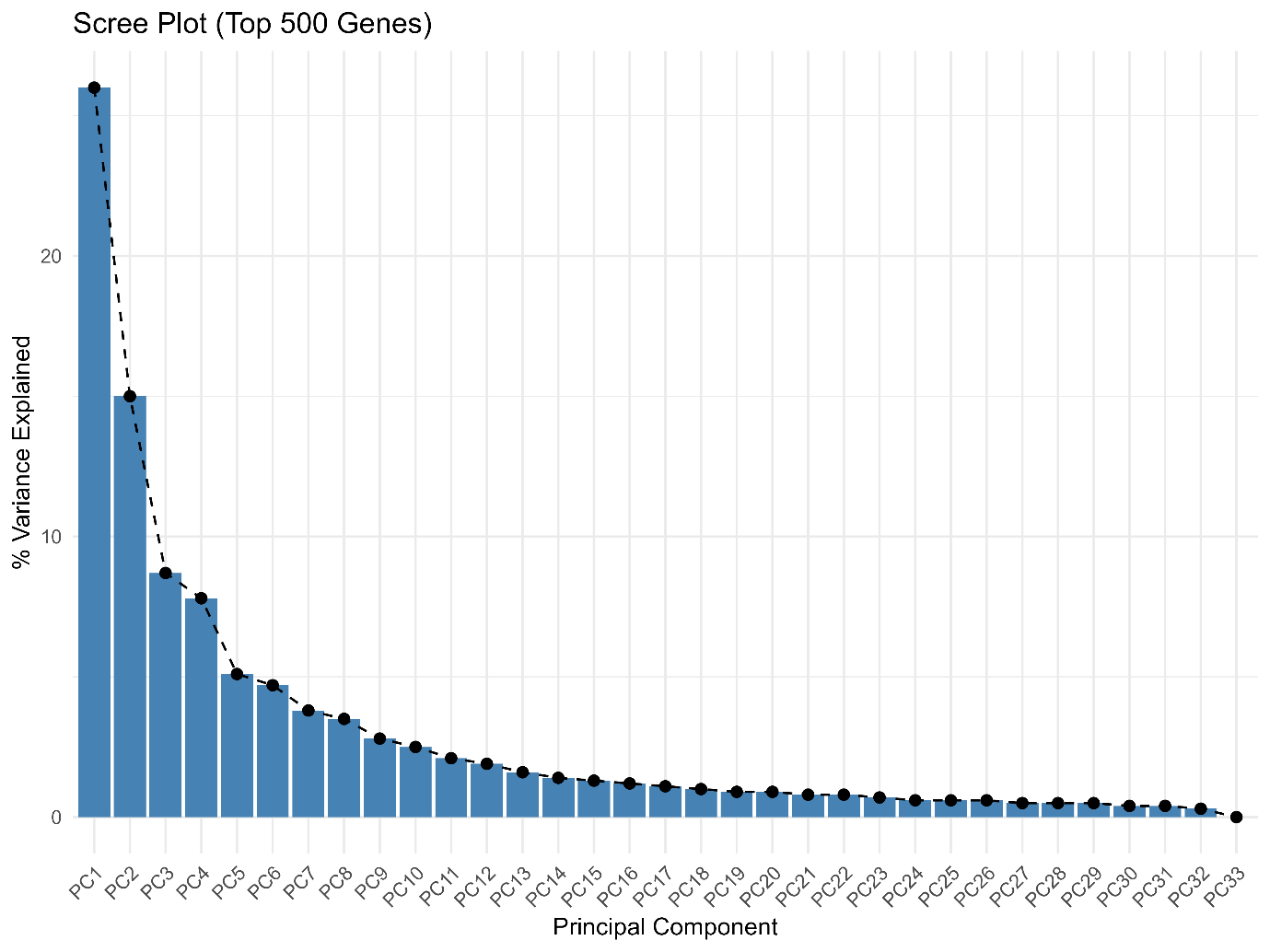

**Figure S2**

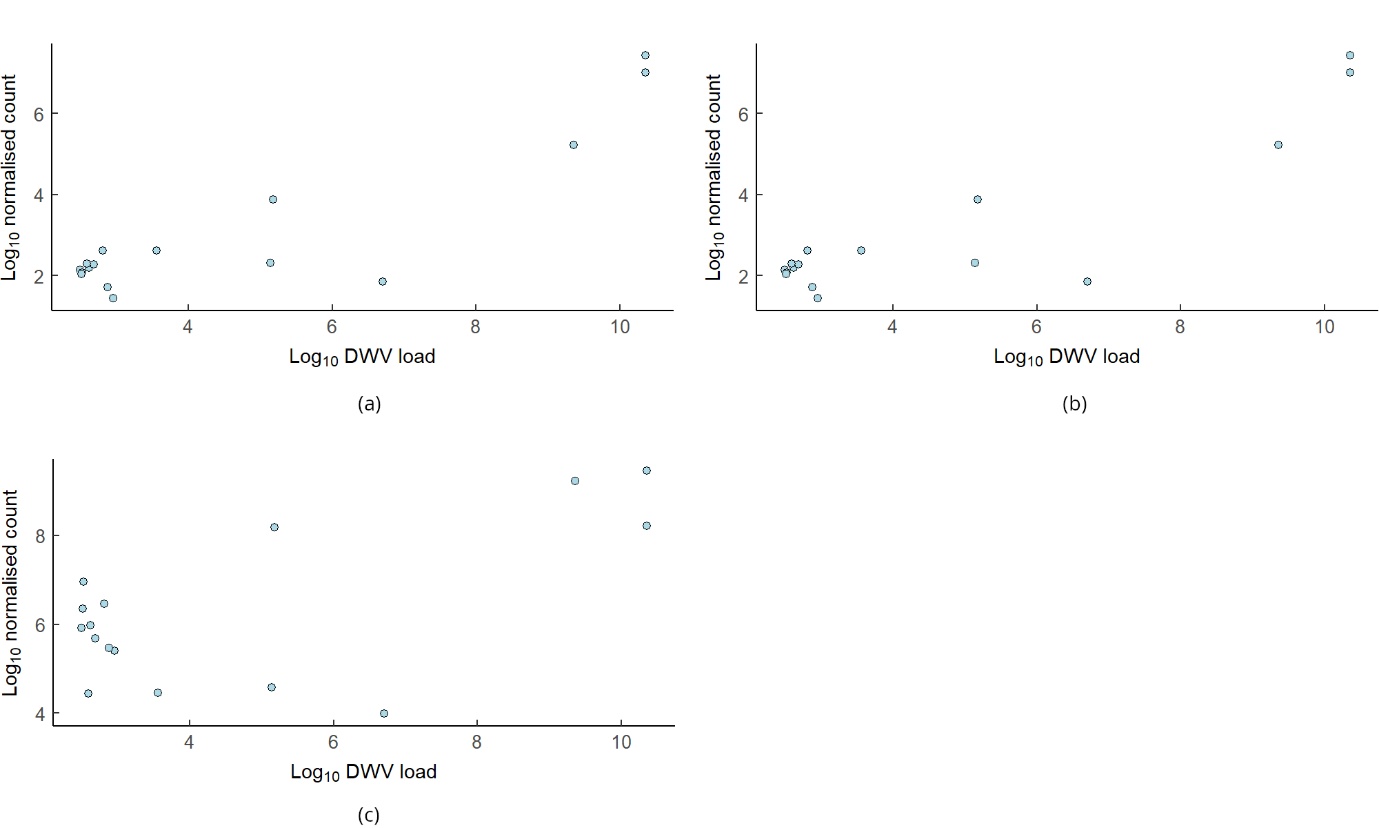

**Figure S3**

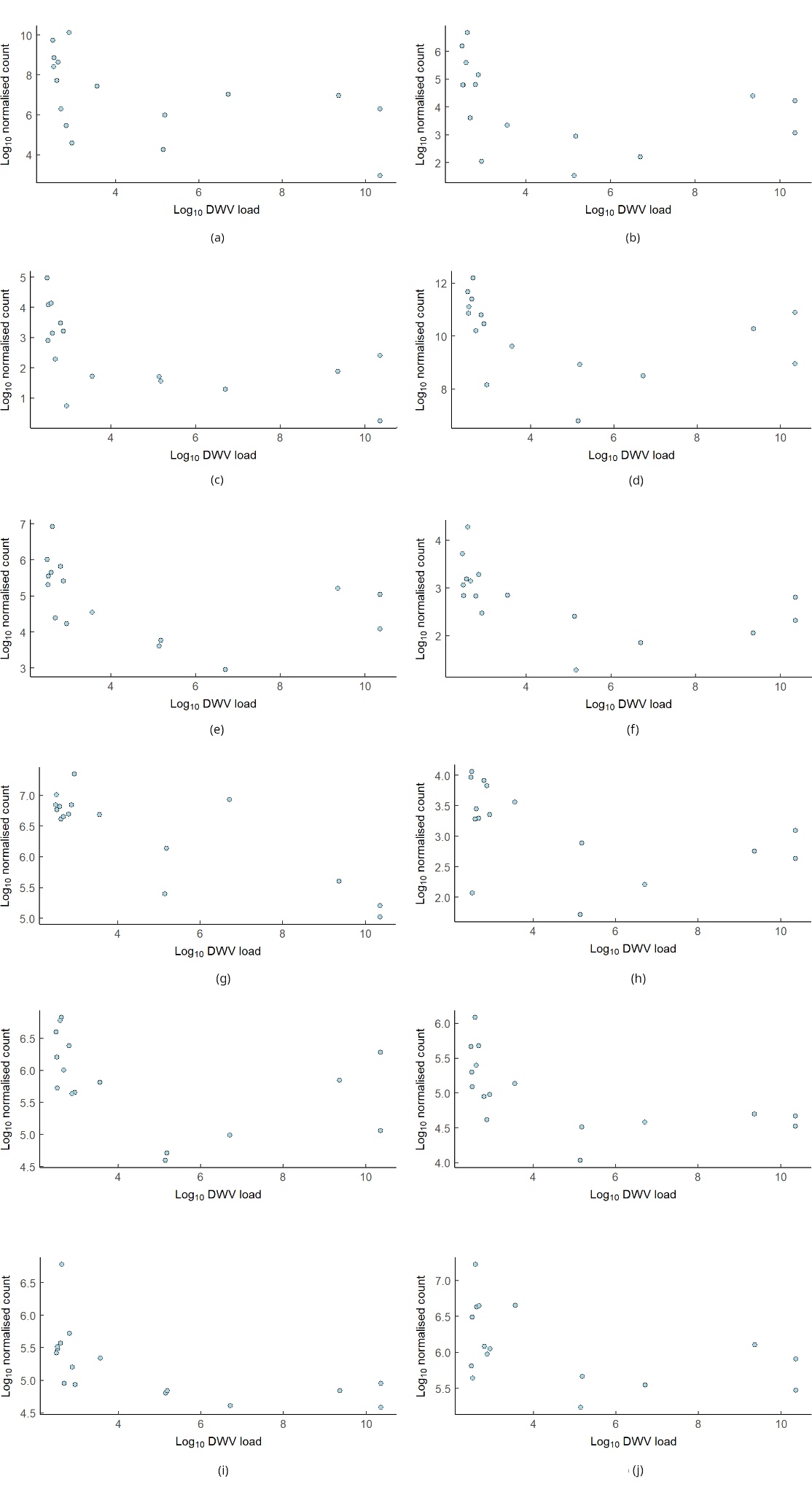

**Figure S4**

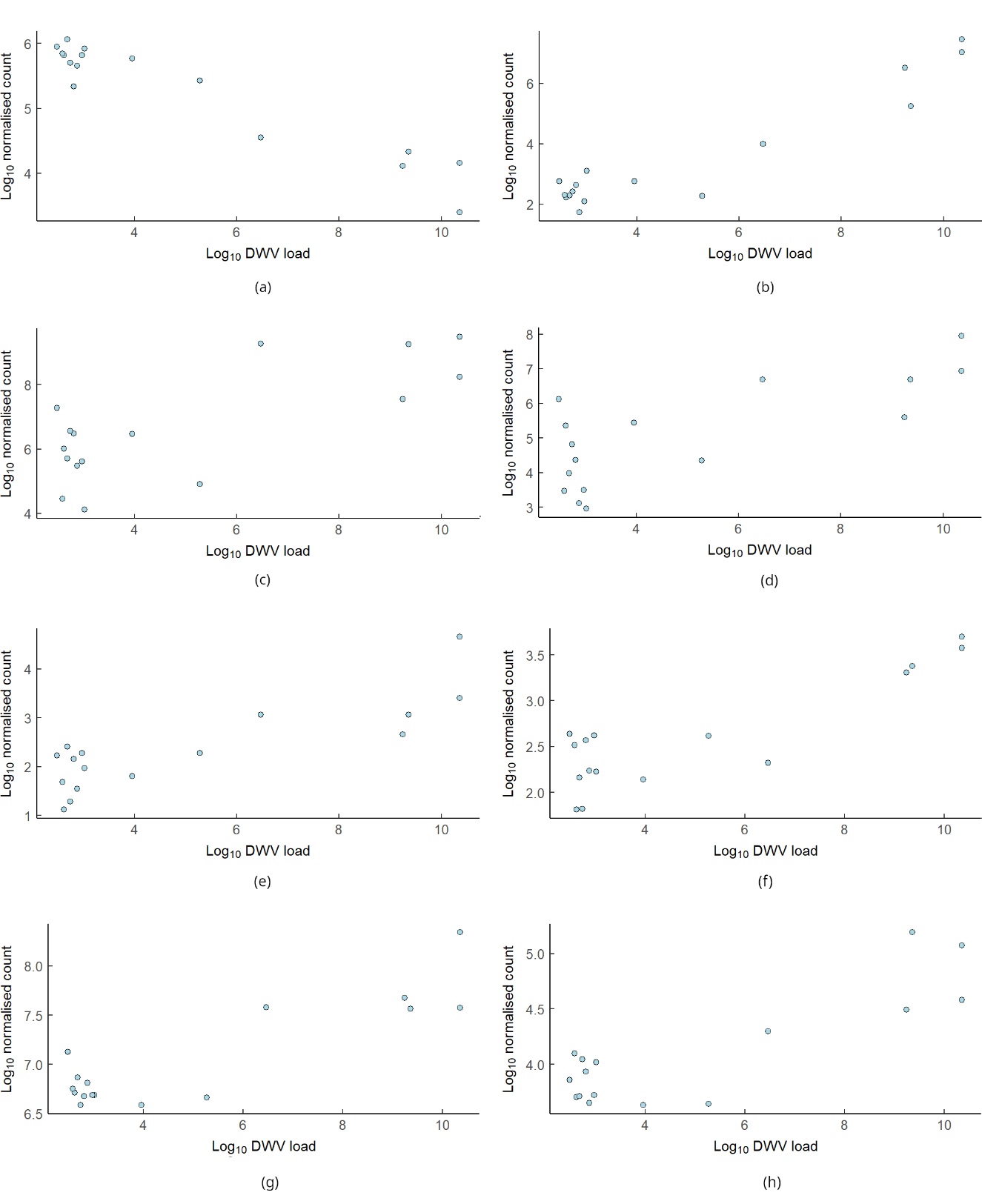

**Figure S5**

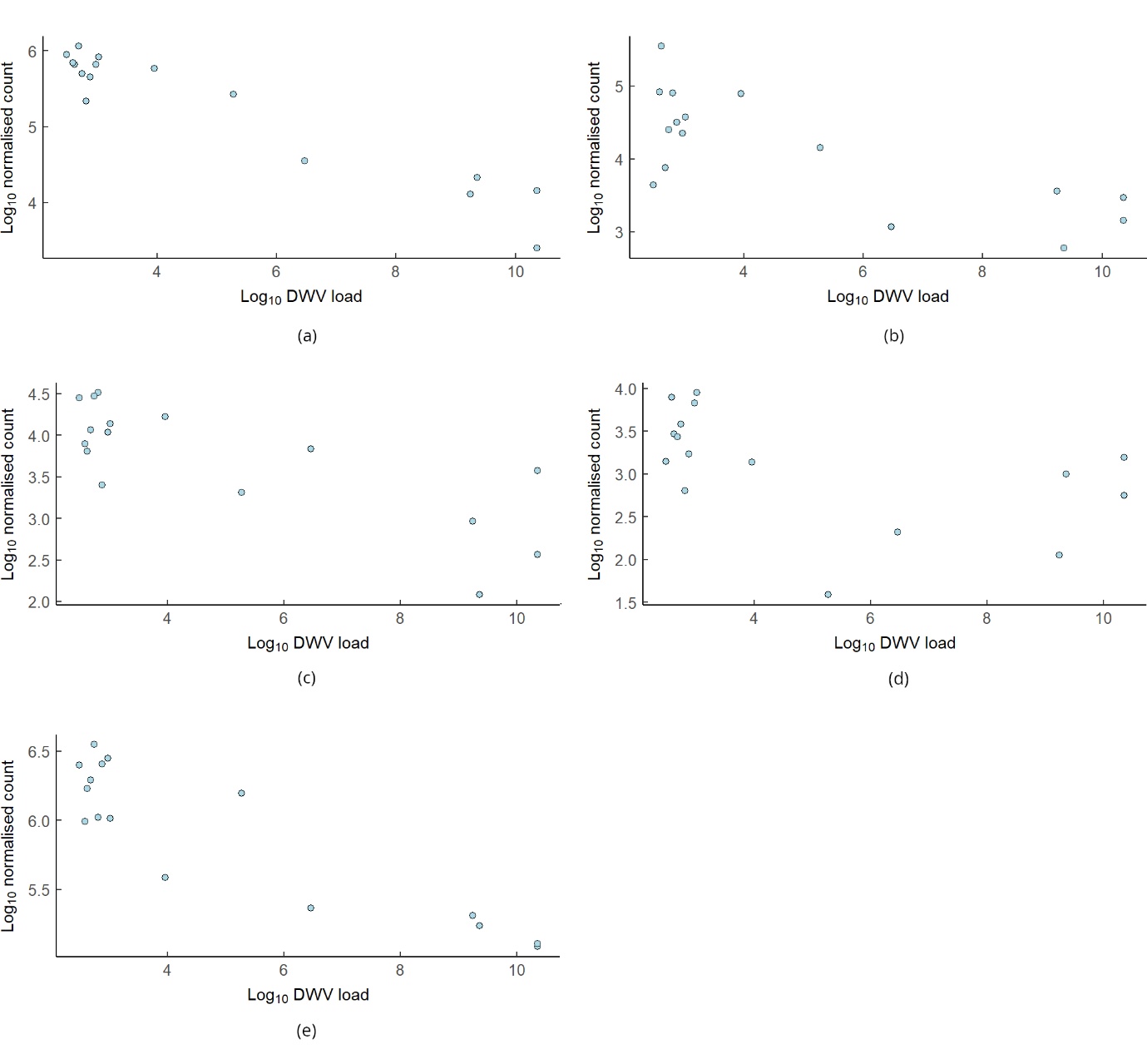

**Table S1**: KEGG pathways identified via overrepresentation analysis of Poor Learners. None of the pathways was significant following a Bonferroni-Hochberg adjustment.

| Regulation ^1^ | KEGG ID | KEGG Description | Genes in Category ^2^ | Differentially Expressed Genes ^3^ |
| --- | --- | --- | --- | --- |
| Up | ame04745 | Phototransduction | 9 | 2 |
|  | ame01200 | Carbon metabolism | 42 | 2 |
|  | ame03040 | Spliceosome | 49 | 2 |
| Down | ame00500 | Starch and sucrose metabolism | 8 | 2 |
|  | ame00981 | Insect hormone biosynthesis | 11 | 2 |
|  | ame00030 | Pentose phosphate pathway | 13 | 2 |

^1^ Up = more highly expressed in Poor Learners with high viral loads; Down = lower expression in Poor Learners with high viral loads. ^2^ The total number of genes in the pathway. ^3^ The number of differentially expressed among those in the pathway.

**Table S2**: KEGG pathways identified via overrepresentation analysis of Colony A bees. None of the pathways was significant following a Bonferroni-Hochberg adjustment.

| Regulation ^1^ | KEGG ID | KEGG Description | Genes in Category ^2^ | Differentially Expressed Genes ^3^ |
| --- | --- | --- | --- | --- |
| Up | ame04130 | SNARE interactions in vesicular transport | 17 | 1 |
|  | ame04624 | Toll and Imd signaling pathway | 27 | 1 |
| Down | ame00981 | Insect hormone biosynthesis | 10 | 3 |
|  | ame00310 | Lysine degradation | 19 | 3 |
|  | ame00380 | Tryptophan metabolism | 9 | 2 |
|  | ame00500 | Starch and sucrose metabolism | 9 | 2 |

^1^ Up = more highly expressed in Poor Learners with high viral loads; Down = lower expression in Poor Learners with high viral loads. ^2^ The total number of genes in the pathway. ^3^ The number of differentially expressed among those in the pathway.

**Table S3**. Genes identified as potential targets of lncRNAs 102655375 in the Poor Learner group and as potential targets of lncRNAs 102654076, 113218947, and 107965291 in the Colony A group

|  |  | **Targets** | | | |
| --- | --- | --- | --- | --- | --- |
| **Group** | **Regulator ^1^** | **OGSv3.2 ID ^2^** | **OGSv1.x ID ^3^** | **Gene ID ^4^** | **Gene Description** |
| Poor Learners | 102655375 | GB49322 | GB14861 | 552711 | anoctamin-4 |
|  |  | GB54313 | GB13155 | 413386 | uncharacterised protein coding |
|  |  | GB54901, GB55397 | GB13035 | 550961, *Vamp7* | vesicle-associated membrane protein 7 |
|  |  | GB41232 | GB14096 | 551949 | ovarian-specific serine/threonine-protein kinase Lok |
|  |  | GB52719 | GB11654 | 726124 | segmentation protein Runt |
|  |  | GB45874, GB45875 | GB18913 | 724760 | G-protein coupled receptor Mth2 |
|  |  | GB42418 | GB15287 | 412372 | pyruvate dehydrogenase phosphatase regulatory subunit, mitochondrial |
|  |  | GB42552, GB42553 | GB19513 | 100577098 | enolase-phosphatase E1 |
|  |  | GB40064 | GB12206 | 102655054 | protein tyrosine phosphatase domain-containing protein 1 |
|  |  |  |  | 102655080 | long non-coding RNA |
|  |  |  |  | 104796166, *Mir9878* | microRNA 9878 |
| Colony A | 102654076 | GB42828 | GB14997 | 409085 | guanine nucleotide exchange factor for Rab-3A |
|  |  | GB55424, GB55425 | GB18331, GB30192 | 725827, *InR-2* | insulin-like receptor-like |
|  |  | GB40969 |  | 102655814 | ras-related and estrogen-regulated growth inhibitor |
|  |  | GB40686 | GB19796 | 551426 | uncharacterised protein coding |
|  | 113218947 | GB45987 |  | 100578618 | uncharacterised protein coding |
|  | 107965291 | GB54319 | GB13586 | 410052, Syt20 | synaptotagmin 20 |
|  |  | GB49625, GB51241 | GB11293 | 100578126 | zinc finger protein 26-like |

^1^ The lncRNA purported to be serving as a regulator. ^2^ Gene IDs as specified in Official Gene Set (OGS) version 3. ^3^ Gene IDs as specified in Official Gene Set (OGS) version 1.x. ^4^ NCBI Gene ID and name (if available) (https://www.ncbi.nlm.nih.gov/gene/
